# A tRNA-derived second messenger mediates antiviral defense

**DOI:** 10.64898/2026.08.25.746378

**Authors:** Yang Liu, Yan Qin, Imane Bouzit, Ann Yeung, Heather Chang, Jonathan Strecker

## Abstract

CRISPR-Cas systems are RNA-guided nucleases that enable prokaryotic immunity; however, some loci encode additional associated genes that cooperate with CRISPR effectors to perform diverse biological functions. Here, we uncover a CRISPR-associated kinase (CASK) system that links the recognition of target RNA to protein phosphorylation. We show that the kinase Csx33 phosphorylates Csx34 following activation of Cas13, enabling Csx34 to bind DNA in a sequence-specific manner. Together, Csx33 and Csx34 function as a transcriptional activation module that upregulates cas13 and associated genes, revealing a positive autoregulatory circuit that potentiates the immune response upon detection of foreign RNA. At the molecular level, Csx33 is activated by CCA trinucleotide RNA generated by Cas13-mediated cleavage of tRNA 3′ tails, uncovering a novel linear second messenger in bacterial immunity and a previously unrecognized signaling role of collateral RNA fragments. Together, these findings establish CASK systems as a new platform for RNA sensing and for engineering programmable phosphorylation-based signaling systems.

## Introduction

Prokaryotes possess numerous defense strategies against foreign genetic elements including CRISPR-Cas systems which typically provide adaptive immunity through a variety of RNA- guided nucleases^1,2^. However, in addition to simple endonuclease activity, bioinformatic searches have identified rare but conserved genes in proximity to CRISPR loci^3–5^ and previous work investigating these genetic associations has uncovered new RNA-guided functions including CRISPR-associated transposases that perform RNA-targeted DNA insertion^6–8^, and CRISPR-associated proteases that cleave protein substrates upon target RNA detection^9–11^. These systems reveal how diverse enzymatic activities can acquire, or be acquired by, RNA-guided effectors and likely only scratch the surface of the programmable functions that exist in nature. Engineering of these natural systems has yielded powerful approaches for genome editing^12^ and RNA sensing in cells^9,13,14^, underscoring the value of continued exploration of microbial diversity.

Distinct classes of CRISPR-associated proteases have been shown to cleave sigma factor inhibitors, thereby linking nucleic acid detection to transcriptional regulation^9,11^. These systems provide a mechanism to coordinate the antiviral response, including the upregulation of spacer acquisition, raising the possibility that detection of foreign genetic elements triggers broader transcriptional programs in their native hosts. Whether CRISPR-Cas systems are integrated with additional signal transduction pathways remains largely unknown. Here, we determine the function and mechanism of a CRISPR-associated kinase (CASK) system that couples RNA detection to protein phosphorylation to regulate the immune response.

## Results

### The CRISPR-associated kinase Csx33 phosphorylates Csx34

Bioinformatic searches for genes associated with CRISPR-Cas systems have identified numerous uncharacterized hits which are prime candidates for enzymes regulated in response to nucleic acid recognition. While searching for genes associated with the RNA-guided RNA nuclease Cas13, we identified a conserved operon containing three genes, here referred to as *csx32, csx33,* and *csx34* (Fig. 1a). Phylogenetic analysis places the Cas13 proteins within the type VI-A family, although not strictly in a monophyletic clade (Fig. 1b and Extended Data Fig. 1a). Csx33 contains an N-terminal kinase domain and a C-terminal tetratricopeptide repeat (TPR) domain, and protein homology analysis reveals similarity to *Mycobacterium tuberculosis* PknG—a eukaryotic-like serine/threonine protein kinase involved in nutrient sensing and virulence^15,16^. The second protein, Csx32, contains a domain of unknown function 3800 (DUF3800), while the third protein, Csx34, has no clear sequence homology to known proteins but exhibits structural similarity to helix–turn– helix (HTH) domains (Fig. 1c).

**Figure 1.**
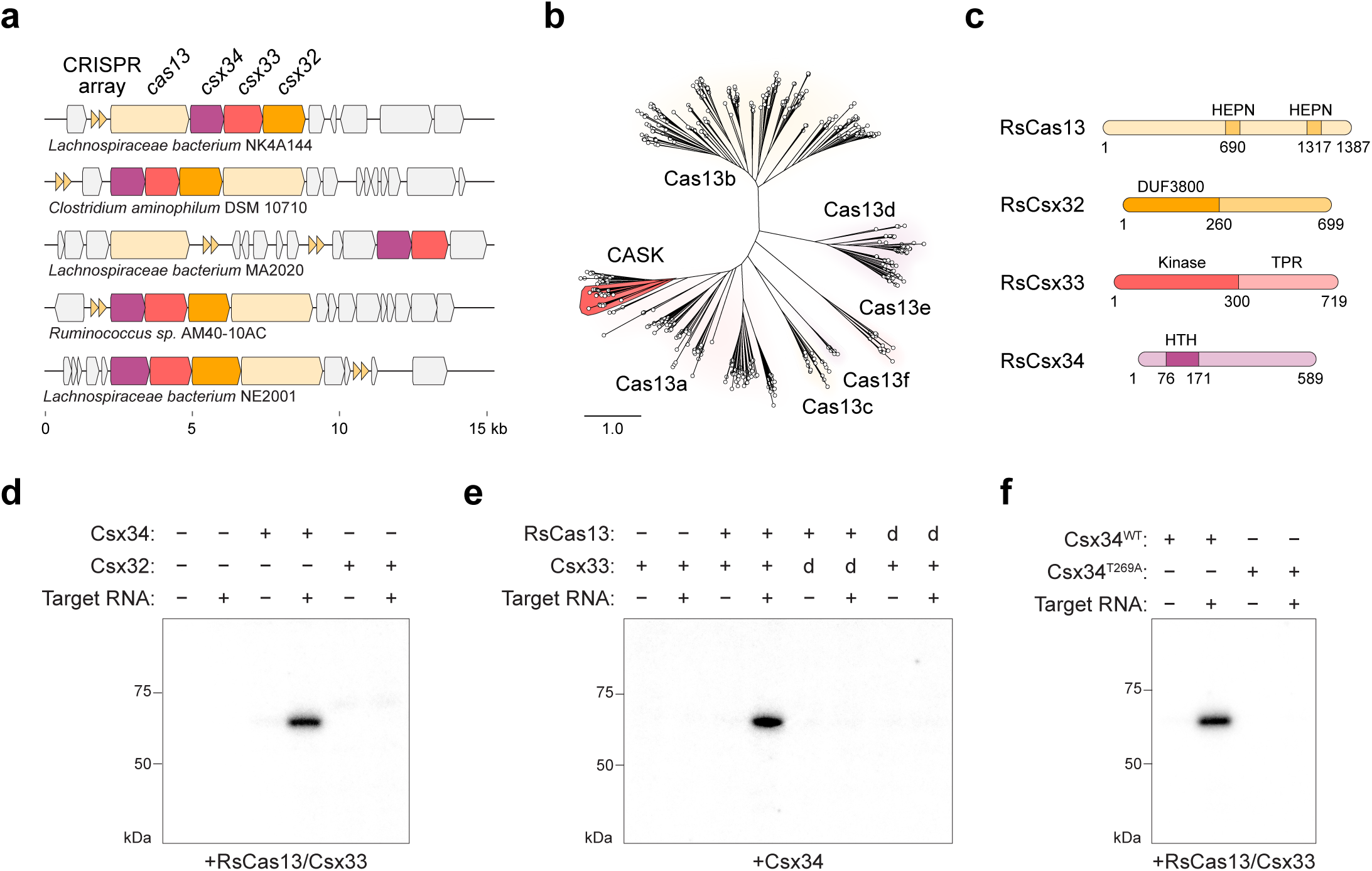
Discovery and reconstitution of a Cas13-associated protein kinase. **a**, Schematic of CASK loci containing three conserved genes in bacteria. **b,** Tree of Cas13 nucleases; CASK systems are highlighted in red. **c,** Domain architecture of CASK proteins. **d,** Csx33 phosphorylates Csx34 in response to target RNA. **e,** Phosphorylation of Csx34 requires the catalytic activity of RsCas13 and Csx33. dRsCas13 (RsCas13:R675A/R1272A), dCsx33 (Csx33:D154A). **f,** Functional validation of the Csx34 phosphorylation site. (d–f) are autoradiographs of SDS–PAGE-resolved in vitro kinase reactions.

Previous studies of CRISPR-associated proteases revealed that their protein substrates are encoded within the same operon, leading us to hypothesize that the kinase Csx33 might similarly phosphorylate one of the neighboring gene products following Cas13 activation. To investigate CASK systems, we synthesized the genes from five identified operons, purified the encoded protein components from *E. coli*, and focused on the system from *Ruminococcus sp*. (RsCASK) because of its conserved kinase catalytic residues and the favorable biochemical properties of its components (Extended Data Figs. 1 and 2).

**Figure 2.**
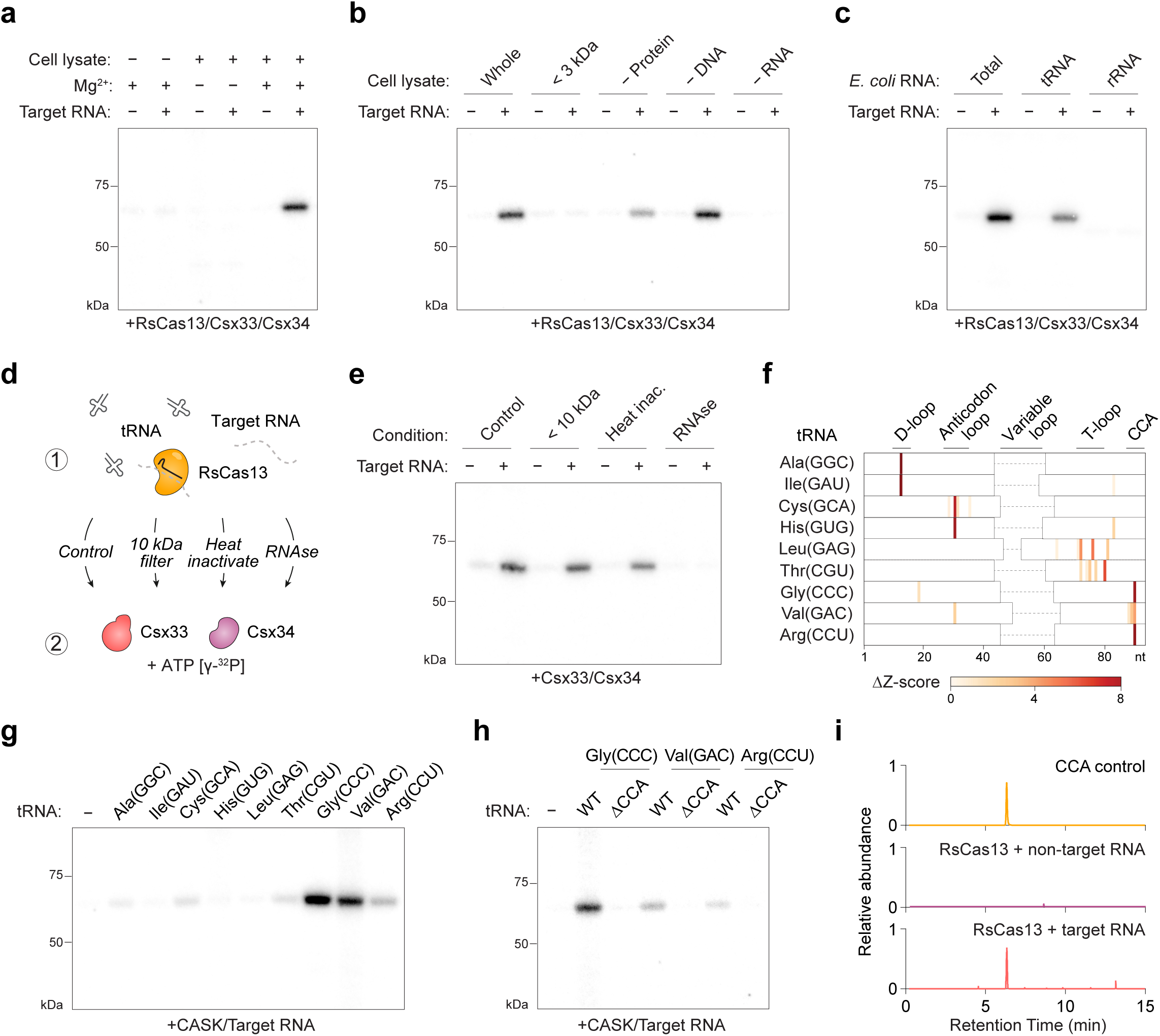
Identification of a cell lysate factor required for Csx33 activity. **a**, CASK reactions require magnesium and *E. coli* cell lysate. **b,** Fractionation of cell lysate uncovers RNA as a necessary component. **c,** Identification of tRNA as the critical factor for CASK reactions. **d,** Schematic of two-pot CASK reactions separating RsCas13 and Csx33. **e,** RsCas13 RNA cleavage products trigger CASK activity. (a–c), (e) and (f) are autoradiographs of SDS–PAGE-resolved in vitro kinase reactions. **f,** RsCas13 collateral cleavage of in vitro–transcribed *E. coli* tRNA. RNA sequencing identifies novel 3′ tRNA ends following in vitro reactions containing target RNA, compared to a non-target RNA control. **g,** In vitro CASK reactions with individual in vitro–transcribed *E. coli* tRNA. **h,** tRNA CCA tails are necessary for CASK activity. **i,** Detection of CCA RNA fragments from RsCas13 collateral reactions containing tRNA^Gly(CCC)^ by liquid chromatograph–mass spectrometry (LC-MS). (a–c), (e), (g), and (h) are autoradiographs of SDS– PAGE-resolved in vitro kinase reactions.

We performed in vitro kinase reactions containing RsCas13, crRNA, Csx33, γ-^32^P-ATP, and the two candidate substrates, and observed radiolabeling of Csx34 but not Csx32 (Fig. 1d). Importantly, this activity was dependent on the addition of target RNA complementary to the crRNA, indicating that Csx33 is activated downstream of RsCas13 upon RNA detection. We were unable to detect autophosphorylation of Csx33 or modification of RsCas13 in reactions. We determined that phosphorylation requires the catalytic activity of the RsCas13 HEPN (higher eukaryotes and prokaryotes nucleotide-binding) nuclease domain and the catalytic loop of Csx33 (Fig. 1e). Mutations in the Csx33 activation segment responsible for ATP binding also impaired activity (Extended Data Fig. 3a). We performed phosphoproteomic analysis of Csx34 following in vitro reactions and observed a single modification on Thr269. Mutation of this residue to alanine (Csx34 T269A) abolished activity in our assay, confirming Thr269 as the sole phosphorylation site (Fig. 1f). Together, our results indicate that Csx33 phosphorylates Csx34 in response to target RNA cleavage by RsCas13 and we next sought to understand the mechanism of kinase activation.

### CASK activity requires tRNA

The C-terminus of Csx33 contains a TPR domain, a motif typically involved in protein–protein interactions, including in the CRISPR-associated protease Csx29, which utilizes this domain to bind Cas7-11^9,10^. We therefore tested whether Csx33 might similarly interact with RsCas13, but did not detect a complex in co-purification experiments. During reconstitution of CASK reactions, we determined that two factors were essential for activity: magnesium and *E. coli* cell lysate (Fig. 2a). The requirement for cell lysate was unexpected, and to test whether small molecules or cofactors were responsible, we filtered lysates through a 3-kDa cutoff membrane, but found that the flow-through did not support activity (Fig. 2b). We next depleted lysates using proteinase K, DNAse I, and immobilized RNase A and found that while removal of proteins and DNA had no effect, the loss of cellular RNA abolished Csx34 phosphorylation in our assay (Fig. 2b). Consistent with this result, addition of purified *E. coli* total RNA supported CASK activity in the absence of cell lysate, which could be traced specifically to tRNA (Fig. 2c).

Following target RNA binding, Cas13 exhibits collateral cleavage of cellular RNAs, including tRNA anticodon loops^17^, leading us to hypothesize that RsCas13-generated tRNA products might activate Csx33. To test this, we performed two-pot CASK reactions in which RsCas13 and RNA components were incubated separately (Reaction 1) before transfer to kinase reactions containing Csx33 and Csx34 (Reaction 2; Fig. 2d). We observed that kinase activity was retained when Reaction 1 was filtered or heat-inactivated, indicating that activation does not require RsCas13 in Reaction 2, but is rather mediated by a small and heat-stable factor (Fig. 2e). Conversely, RNase A treatment abolished the ability of Reaction 1 to activate Csx33, supporting a model in which the activation signal is a cleaved tRNA product (Fig. 2e). We investigated whether diverse Cas13 members might also activate Csx33 through collateral tRNA cleavage, or if RsCas13 possesses unique properties. We tested three additional Cas13 nucleases: LwaCas13a, LbuCas13a, and RfxCas13d^18,19^, and confirmed cleavage of target RNA, but were unable to detect Csx34 phosphorylation in kinase reactions (Extended Data Fig. 3b–d).

### RsCas13 collateral activity generates diverse tRNA fragments

To identify the tRNA fragments responsible for Csx33 activity, we first performed RNA sequencing of RsCas13 collateral digests of in vitro–transcribed *E. coli* tRNA. Mapping of unique 3′ RNA ends revealed cleavage positions in D-loops, anticodon loops, T-loops, and 3′ CCA tails (Fig. 2f and Extended Data Fig. 4). Testing individual in vitro–transcribed tRNAs revealed that CASK activity strongly correlates with 3′-tail cleavage, including for tRNA^Gly(CCC)^ and tRNA^Val(GAC)^ (Fig. 2g). Sequence analysis of cleaved tRNA 3′ tails revealed no nucleotide preference at the −1 position upstream of the cleavage site, whereas other tRNA fragments showed only minor enrichment for uridine at this position (Extended Fig. 5a).

To directly test whether the 3′ CCA sequence is required for Csx33 activity, we compared three wild-type tRNAs with their corresponding truncated variants (ΔCCA) and found that loss of CCA sequence abolished CASK activity (Fig. 2h). We confirmed RsCas13-mediated collateral cleavage of tRNA^Gly(CCC)^ by denaturing gel electrophoresis, and systematic testing of different tRNA segments also revealed that the 3′ end is necessary and sufficient for CASK activity (Extended Data Fig. 5b,c). Short RNAs corresponding to the final 10 nucleotides of certain tRNAs also supported CASK activity and required the presence of the 3′ CCA tail (Extended Data Fig. 5d).

Due to the small size of the CCA fragments, our RNA sequencing method—which utilizes silica-based purification columns—captured only the larger 5′ fragment of these cleavage events. To directly test whether RsCas13 generates CCA trinucleotides, we analyzed collateral cleavage reactions containing tRNA^Gly(CCC)^ by liquid chromatography–mass spectrometry (LC–MS). This analysis revealed target RNA–dependent generation of CCA fragments, which matched the retention time and mass-to-charge ratio of a CCA positive control (Fig. 2i and Extended Data Fig. 6a,b).

### Csx33 is activated by CCA trinucleotide RNA

Together, these results reveal that RsCas13 cleaves tRNA 3′ tails to generate a CCA trinucleotide RNA, hereafter referred to as CCA RNA, and that the CCA sequence is required for CASK activity. We next tested whether CCA RNA is sufficient to activate Csx33. Although short RNAs derived from the 3′ end of tRNA^Gly(CCC)^ supported CASK activity, in vitro–transcribed CCA RNA unexpectedly did not (Fig. 3a), suggesting that Csx33 activation might also depend on the chemical ends generated by RsCas13 cleavage. In vitro transcription by T7 RNA polymerase produces RNAs with 5′-triphosphate ends, whereas cleavage by Cas13 and other metal-independent RNases generates products with 2′,3′-cyclic phosphate and 5′-OH ends. To assess the importance of the 5′ phosphorylation state, we treated 5′-ppp–CCA-3′-OH RNA with RNA 5′ pyrophosphohydrolase (RppH) to generate a 5′ monophosphate RNA, or calf intestinal alkaline phosphatase (CIP), to remove all 5′ phosphates (Fig. 3b). We observed that CIP treatment, but not RppH treatment, enabled CCA RNA to activate Csx33 in the absence of Cas13 (Fig. 3c), indicating that a 5′-OH end is required for CASK activity. In an orthogonal approach, we generated a second CCA RNA containing a 5′ poly-G extension and treated it with RNase T1, which cleaves after guanosine residues to yield a CCA fragment bearing a 5′-OH end (Fig. 3b). RNase T1-treated CCA RNA similarly activated Csx33, and this activity was abolished by subsequent phosphorylation with T4 polynucleotide kinase (PNK), confirming that the absence of a 5′ phosphate is critical for Csx33 activation (Fig. 3c).

**Figure 3.**
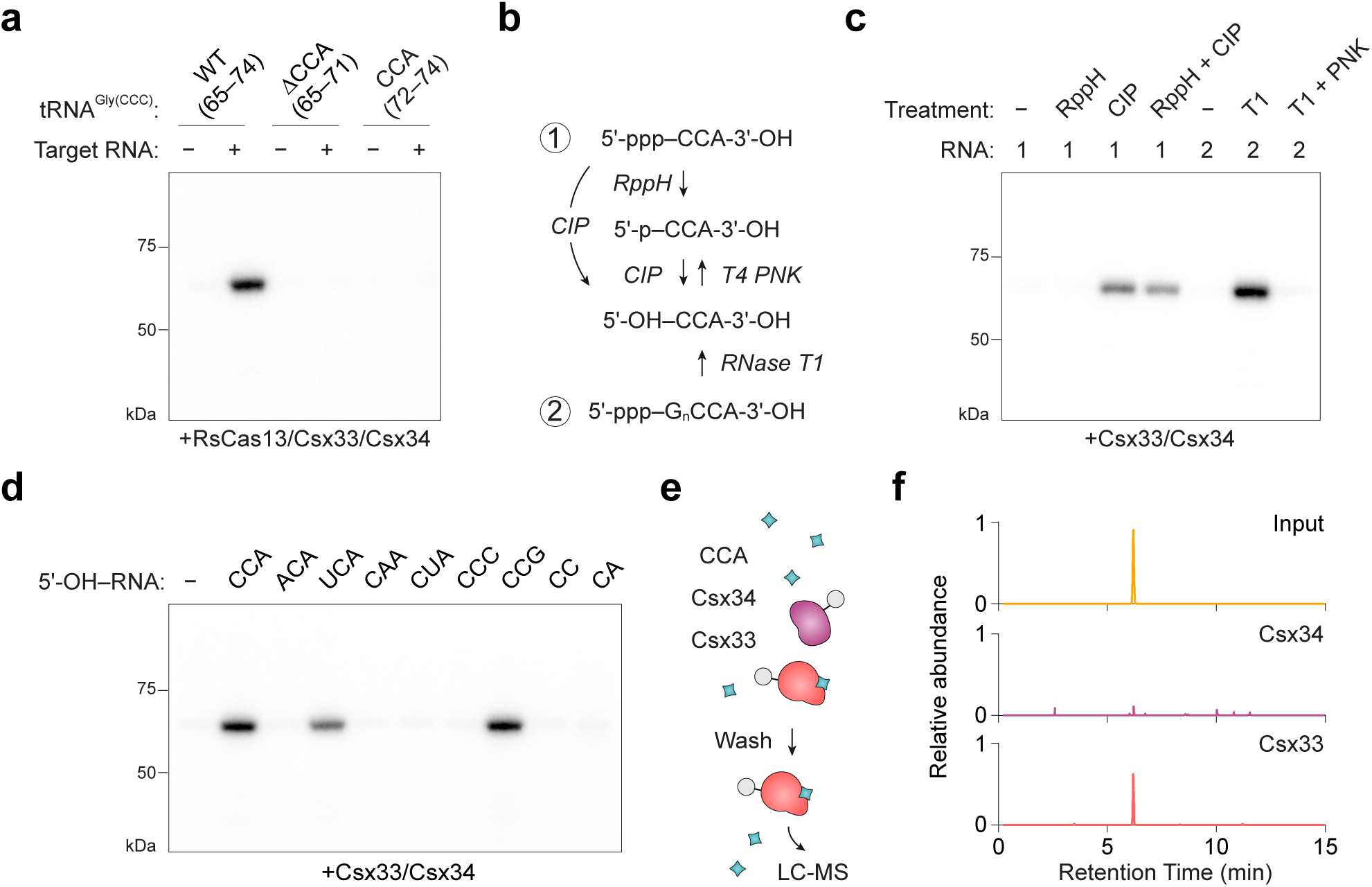
Csx33 is activated by a CCA trinucleotide RNA. **a**, CASK activity requires a CCA-containing RNA and Cas13 cleavage. **b,** Schematic of reactions to produce 5′-OH ends from in vitro–transcribed RNA using RNA 5’ pyrophosphohydrolase (RppH), calf intestinal alkaline phosphatase (CIP), RNase T1 (T1), and T4 polynucleotide kinase (PNK). **c,** Csx33 is activated by 5′-OH–CCA. **d,** Specificity of Csx33 for different oligonucleotide sequences. **e,** Experimental design to detect CCA binding to immobilized CASK proteins. **f,** Csx33 binds 5′-OH–CCA RNA, as detected by LC– MS. (a), (c), (d) and (f) are autoradiographs of SDS–PAGE-resolved in vitro kinase reactions.

We tested the specificity of Csx33 for different RNA fragments and found that, in addition to CCA, Csx33 could be triggered by UCA and CCG trinucleotides containing single transition mutations, whereas transversion mutations and changes at the central cytosine position were not tolerated (Fig. 3d). We did not detect phosphorylation with dinucleotide RNAs, and tetranucleotide RNAs showed greatly reduced activity, revealing that Csx33 preferentially responds to trinucleotide RNA (Fig. 3d and Extended Data Fig. 6c). Independent of Cas13, we observed that Csx33 also requires magnesium for activity (Extended Data Fig. 6d). To test whether Csx33 directly senses CCA RNA, we immobilized CASK proteins, incubated them with 5′-OH–CCA, and analyzed bound RNA by LC–MS (Fig. 3e). Notably, we detected CCA bound to Csx33, but not to Csx34 (Fig. 3f).

### Csx34 phosphorylation enables DNA binding

Having identified the activation mechanism, we next investigated the biological function of Csx34 and how it is regulated by phosphorylation. We first tested whether Csx33 and Csx34 contribute to Cas13-mediated defense against a plasmid-encoded transcript in *E. coli*; however, their addition had no effect in this assay (Extended Data Fig. 7). Csx34 contains a predicted HTH domain (residues 76–171), a fold commonly found in DNA-binding proteins and transcriptional regulators^20^, and we therefore hypothesized that phosphorylated Csx34 might bind DNA. To test this and identify putative binding motifs, we performed systematic evolution of ligands by exponential enrichment (SELEX) using immobilized wild-type Csx34 and a phosphomimetic Csx34 T269E mutant (Fig. 4a). We detected a DNA motif with dyad symmetry that was progressively enriched over successive rounds of selection and was specific to the Csx34 T269E- bound samples (Fig. 4b,c and Extended Data Fig. 8). Fluorescence anisotropy assays confirmed that Csx34 T269E binds DNA containing the SELEX-derived motif, but not a random control sequence, and that binding requires the phosphomimetic mutation (Fig. 4d,e). To test the ability of natively phosphorylated Csx34 to bind DNA using an orthogonal assay, we performed in vitro CASK reactions with tRNA and 300-bp DNA substrates and analyzed complex formation by mass photometry. We detected a larger DNA–protein complex that required both the Csx34-binding motif and target RNA in the reaction (Fig. 4f) and mirrored a Csx34 T269E–DNA complex (Extended Data Fig. 9).

**Figure 4.**
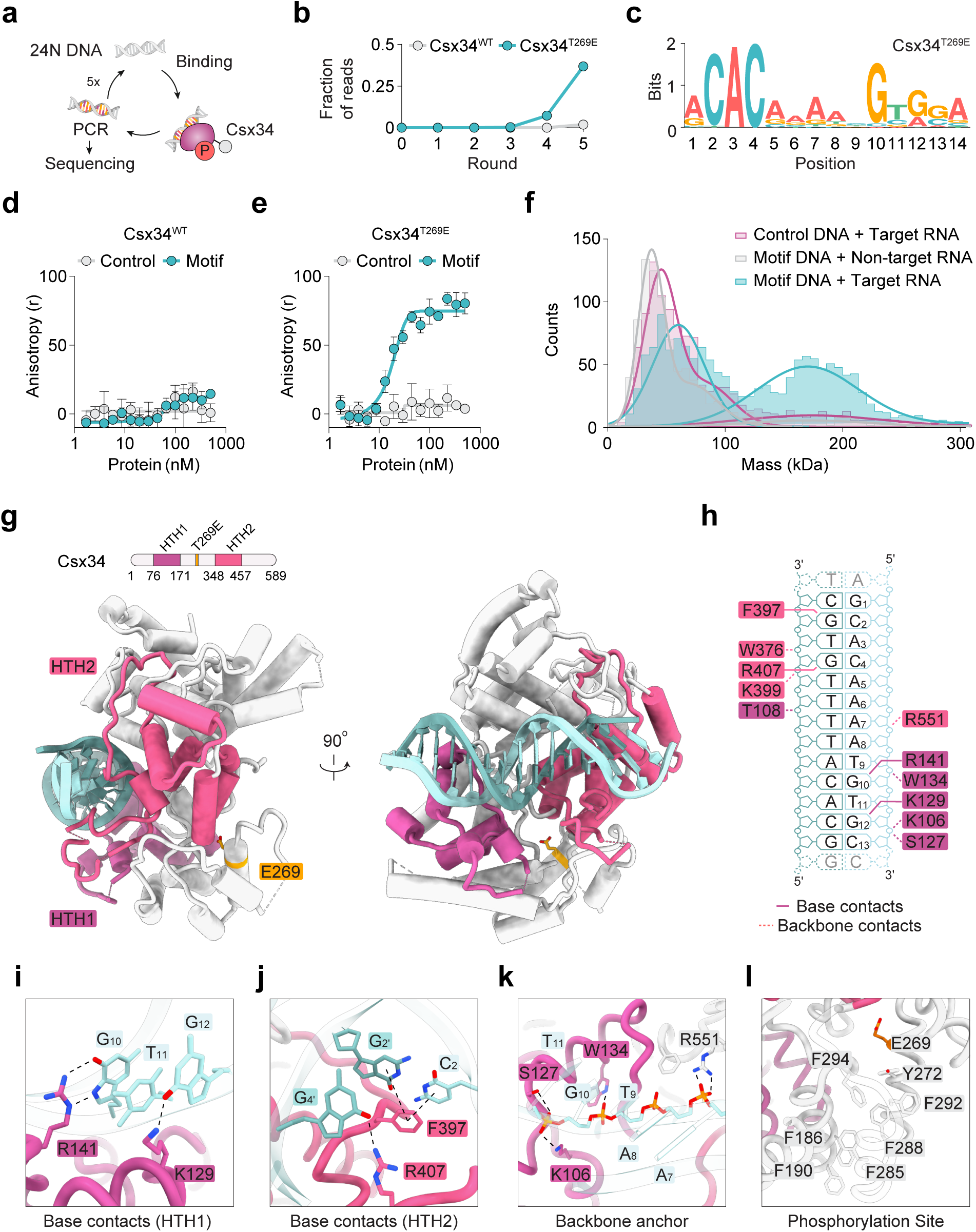
Phosphorylation of Csx34 enables DNA binding. **a**, Schematic of Systematic Evolution of Ligands by Exponential Enrichment (SELEX) experiments. **b,** Enrichment of a Csx34 T269E binding motif over successive rounds of SELEX, plotted as a fraction of the total sequenced reads. **c,** Sequence of the identified Csx34 T269E DNA binding motif. **d, e,** Binding of wild- type Csx34 (d) and Csx34 T269E (e) protein to 15 bp DNA substrates measured by fluorescence anisotropy. **f,** DNA binding of Csx34 following in vitro CASK reactions in the presence of target or non-target RNA, measured by mass photometry. **g,** Cryo-EM structure of Csx34 T269E bound to a 15 bp DNA substrate. The phosphomimetic residue Glu269 is highlighted in orange. **h,** Schematic of the resolved DNA bases and overview of key contacts made by Csx34. **i,** Base contacts of the HTH1 domain. **j,** Base contacts of the HTH2 domain. **k,** Anchoring of the DNA backbone. **l,** Close-up of Glu269 above a hydrophobic pocket between the two HTH domains.

To understand how phosphorylated Csx34 engages DNA, we determined the structure of a Csx34 T269E–DNA complex using cryo-electron microscopy (cryo-EM) at a resolution of 3.1 Å (Fig. 4g and Extended Data Fig. 10a,b). Structural analysis revealed a second HTH domain (residues 348– 457), with each HTH domain binding to a GTG motif within the DNA substrate (Fig. 4h–j and Extended Data Fig. 10c,d). This sequence-specific recognition is driven by a network of base-specific interactions. Within HTH1, residues Arg141 and Lys129 recognize guanines G10 and G12 (Fig. 4i), whereas in HTH2, contacts to guanines G2′ and G4′ are mediated by Phe397 and Arg407 (Fig. 4j). Additional backbone interactions involving residues Lys106, Thr108, Ser127, and Trp134 in HTH1, and Trp376, Lys399, and Arg551 in HTH2, further stabilize DNA binding (Fig. 4h,k). In our structure, the phosphomimetic residue Glu269 is located within an alpha-helical bundle that bridges HTH1 and HTH2, facilitating the formation of a stacked hydrophobic core including residues Phe186, Phe190, Tyr272, Phe285, Phe288, Phe292, and Phe294 (Fig. 4l).

Comparison of our Csx34 T269E–DNA structure with a high-confidence prediction of apo–Csx34 reveals substantial rearrangement of HTH2, while HTH1 and the rest of the protein remain largely unchanged (Extended Data Fig. 11a–d). Notably, Phe397 and Arg407 are repositioned in the Csx34 T269E–DNA structure to recognize two key guanines in the DNA substrate. Together, these data support a model in which phosphorylation regulates DNA binding by remodeling HTH2 to orient critical DNA-interacting residues.

### CASK signaling amplifies the antiviral response

Having defined the Csx34-binding motif, we next searched for potential binding sites in the native host. Although a complete genome sequence for the *Ruminococcus* species harboring the CASK system is unavailable, we identified a prominent match within the 37-kb contig, located between the CRISPR array and the CASK operon (Fig. 5a). To investigate the function of Csx34 and this sequence in bacteria, we constructed an *E. coli* transcriptional reporter by placing 435 bp of native *Ruminococcus* sequence upstream of the *luxCDABE* luciferase operon (Fig. 5b). We observed an upregulation of luciferase activity upon the addition of Csx34 T269E, but not wild-type Csx34, and this effect was abolished by mutating the Csx34-binding motif within the reporter (Fig. 5c). Unexpectedly, robust signal in this assay required co-expression of Csx33, suggesting that both proteins might function together to regulate gene expression, even after phosphorylation (Extended Data Fig. 12a,b). Consistent with this hypothesis, we were able to purify a stable Csx33–Csx34 T269E complex from *E. coli* (Extended Data Fig. 12c–e).

**Figure 5.**
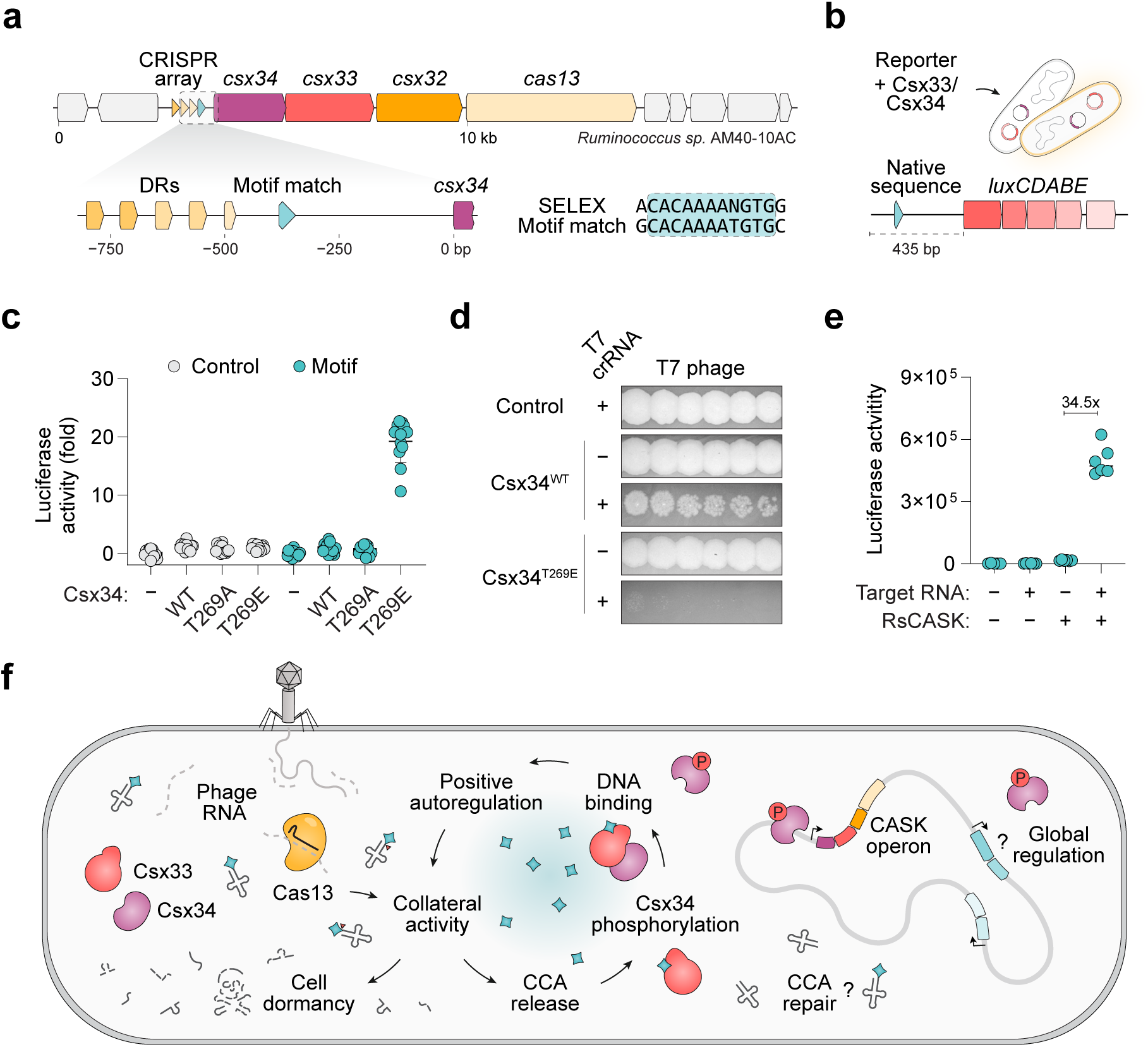
CASK signaling amplifies the antiviral response. **a**, Schematic of the CASK locus from *Ruminococcus sp. AM40-10AC* and a sequence match to the experimentally determined Csx34 binding motif. **b,** Overview of a transcriptional reporter assay in *E. coli* using native *Ruminococcus sp.* sequence. **c,** Csx33–Csx34-mediated transcriptional activation is dependent on Csx34 T269E. Luciferase expression was normalized to cells containing an empty vector. *N* = 12 replicates. **d,** Phage defense by LbuCas13 encoded downstream of the native *Ruminococcus sp.* sequence. T7 phage were diluted 5-fold and spotted on top agar containing *E. coli* strain C expressing Csx33–Csx34 and a T7 targeting (+) or non-targeting (–) crRNA. **e,** Sensing of a programmed target RNA in *E. coli* expressing RsCASK components and a targeting (+) or non-targeting (–) crRNA. *n* = 6 replicates. **f,** Model of CASK activation mechanism and function. In (c) and (e), error bars represent standard deviation from the mean.

We performed tiling mutagenesis of the *Ruminococcus* regulatory sequence and found variants which markedly decreased luciferase activity (R3–R5; Extended Data Fig. 13a,b). Investigation of this region identified a predicted σ^70^-type promoter (P1), and mutation of the –35 and –10 elements similarity reduced basal luciferase activity (Extended Data Fig. 13c,d). Despite this decrease, addition of Csx33–Csx34 T269E exhibited an even higher fold-change in this reporter, compensating for the weakened promoter activity (Extended Data Fig. 13e). To determine whether Csx33–Csx34-mediated transcriptional activation requires host RNA polymerase, we replaced the native promoter with a T7 promoter. Expression of T7 RNA polymerase in these cells resulted in a 100-fold increase in luciferase activity, confirming functionality of our orthogonal system (Extended Data Fig. 13f,g). However, in contrast to its activation of *E. coli* RNA polymerase, Csx33–Csx34 T269E had no effect on T7 RNA polymerase-mediated luciferase output (Extended Data Fig. 13h).

These data reveal that one function of CASK signaling is positive autoregulation of its own operon. To test this functionally, we replaced luciferase in our reporter with LbuCas13 and challenged cells with T7 phage. We observed a partial reduction of phage plaques following expression of wild-type Csx33–Csx34 and a T7-targeting crRNA, but that expression of Csx33–Csx34 T269E greatly enhanced antiviral defense (Fig. 5d).

CASK signaling therefore provides a novel method to activate bacterial gene expression or trigger DNA binding in response to programmed RNA targets. To test whether the full CASK system can be reconstituted in *E. coli*, we co-expressed a target RNA and the three protein components (Csx33, Csx34, and RsCas13) under low induction conditions to minimize RsCas13 collateral toxicity (Extended Data Fig. 14a,b). Addition of a targeting crRNA led to a 34-fold increase in luciferase activity compared to non-targeting controls (Fig. 5e). This signal was completely dependent on Csx34 Thr269, as it was abolished in the T269A mutant, and matched the luciferase signal produced by the constitutively active Csx34 T269E mutant (Extended Data Fig. 14c). Finally, we investigated whether Csx34 could be repurposed as an RNA-regulated scaffold to recruit heterologous effectors to DNA. Fusion of Csx34 to the transcriptional activator SoxS^21^ further increased reporter signal-to-noise ratio, demonstrating that CASK signaling can be used to control synthetic transcriptional circuits (Extended Data Fig. 14d,e).

## Discussion

Here, we discover a link between CRISPR-Cas13 immune systems and the activation of a protein kinase through a tRNA-derived messenger. CASK signaling triggers a transcriptional response, and our data support a model in which phosphorylated Csx34 binds upstream of the promoter as a class-I activator to facilitate host RNA polymerase recruitment. One transcriptional target of this circuit is the CASK operon itself, containing *csx32*, *csx33*, *csx34*, and *cas13*. This reveals a new mechanism of positive autoregulation in CRISPR immunity, directly coupling detection of foreign RNA to potentiation of the antiviral response (Fig. 5f).

CRISPR-Cas systems can impose a high fitness cost on the host^22,23^ creating an evolutionary trade-off between robust immunity and the danger of self-targeting. Consequently, their activity is regulated at multiple levels, including spacer acquisition^9,24–27^, and mechanisms to repress basal activity^28^. CASK signaling provides a distinct form of positive feedback in which Cas13 activity upregulates expression of the CRISPR effector itself. We propose that this regulatory logic allows cells to remain poised for defense, maintaining low basal activity while retaining the capacity for signal-dependent amplification of the immune response. This circuit could also enable fine-tuning of Cas13 expression, thereby modulating collateral cleavage and the induction of cell dormancy according to infection severity. The use of phosphorylation to control transcription is reminiscent of bacterial two-component systems, which couple environmental sensing to response regulator activation and broad transcriptional programs^29^. We hypothesize that Csx33–Csx34 might also upregulate other defense genes throughout the host and future work will be required to characterize the scope of CASK signaling during viral infection.

The 3′ CCA tail is ubiquitous in all mature tRNAs and serves as a critical interface between nucleic acids and protein. Collateral cleavage of tRNA 3′ tails by RsCas13 is therefore expected to impair protein synthesis, consistent with cellular dormancy and tRNA anticodon-loop cleavage associated with other Cas13 nucleases^17,30^. Unexpectedly, our work reveals that tRNA-derived CCA RNA acts as a signaling molecule, establishing a previously unrecognized role for collateral RNA fragments in CRISPR biology. This discovery raises questions about the identity and function of additional effectors that respond to CCA RNA, as well as the broader scope of tRNA-derived messenger signaling in bacterial immunity. Future structural studies will be required to investigate how CCA binding triggers kinase activity and the molecular basis of messenger discrimination. RsCas13 is uniquely able to activate Csx33 in our assays, although we note that other Cas13 nucleases may generate CCA RNA at lower levels (Extended Data Fig. 15). This hints at a previously unrecognized specialization among CRISPR nucleases, in which collateral cleavage products may have evolved to engage specific protein effectors. The observation that Cas12a3 nucleases also cleave tRNA 3′ tails^31^ suggests that CCA-mediated signaling likely extends beyond the CASK system characterized here.

The tRNA-derived CCA fragment therefore establishes a distinct class of linear second messengers in bacterial immunity, expanding the repertoire of nucleotide-based signals beyond cyclic oligoadenylates produced by Cas10 in type III CRISPR-Cas systems^32,33^ and cyclic dinucleotides generated by cGAS-like enzymes^34–36^. A key feature of CCA messengers identified here is the absence of a 5′-phosphate end, which is critical for Csx33 activation. Because bacterial housekeeping and tRNA-processing RNases, such as RNase E and RNase PH, generally produce 5′-phosphate fragments ^37–40^, we propose that the 5′-OH end may serve as a unique signature of Cas13 cleavage and safeguard against activation by routine RNA turnover. Cyclic oligonucleotide messengers are distinguished by their unique linkage chemistry and require dedicated nucleases for degradation, thereby conferring chemical stability. Although linear CCA RNA is likely less stable, its short half-life may be offset by the high cellular abundance of tRNAs and their continual repair by the CCA-adding enzyme, enabling sustained messenger generation.

Finally, this work identifies a new molecular function that can be triggered in response to programmable RNA of interest. Importantly, we demonstrate that CASK systems can be reconstituted as RNA sensors in bacteria without inducing collateral RNA toxicity. We anticipate that future engineering of CASK systems could enable new methods to control protein phosphorylation in specific cell types based on transcriptomic signatures and harness the DNA-binding activity of Csx34 for new purposes in response to detection of specific RNAs. This work provides another example of CRISPR systems coordinating diverse molecular functions beyond nuclease activity, and we expect that the continued exploration of CRISPR-associated enzymes will uncover additional interesting, and potentially useful, RNA-guided systems.

## Methods

### Gene synthesis and cloning

CASK genes were identified by MMseqs2^41^ clustering of Cas13-containing loci. Csx33 was previously identified as a putative Cas13-associated gene but classified as an “unclear” association^3^. All genes were codon-optimized for *E. coli* expression (GenScript) and synthesized as DNA fragments (Azenta). PCRs were performed using Phusion Flash High-Fidelity PCR Master Mix (Thermo Scientific). Cloning was performed by Gibson assembly (NEB), transformed into *E. coli* 5α cells (NEB, C2987), and all plasmids were sequence-verified by in-house Tn5 sequencing.

### In vitro RNA synthesis

Short RNAs (e.g., crRNA and target RNA) were generated by annealing a DNA oligonucleotide containing the reverse complement of the desired RNA with an oligonucleotide encoding a T7 promoter. Standard in vitro transcription reactions were performed using the HiScribe T7 High Yield RNA Synthesis Kit (NEB) at 37 °C for 8–12 h, and RNA was purified using RNAClean XP beads (Beckman Coulter). Full-length tRNA genes were cloned from *E. coli* into pUC19 plasmids containing the hepatitis delta virus (HDV) ribozyme for 3′ processing. These plasmids were amplified by PCR and used as templates for in vitro transcription. tRNAs with processed HDV- generated 3′ ends were repaired with T4 polynucleotide kinase (NEB) and repurified using RNAClean XP beads.

CCA RNA messengers were produced by in vitro transcription for 3 h to limit excessive run-off transcription and purified using RNAClean XP beads (Beckman Coulter). Short RNAs from in vitro reactions were treated with 5′ RNA polyphosphatase (RppH; NEB), calf intestinal alkaline phosphatase (CIP; NEB), RNase T1 (Thermo Scientific), or T4 polynucleotide kinase (NEB) under the manufacturers’ recommended conditions. Reactions were heat-inactivated at 95 °C for 10 min and stored at −80 °C prior to use.

### Protein purification

Genes were cloned into pSUMO plasmids encoding an N-terminal TwinStrep–SUMO tag, and proteins were expressed in *E. coli* BL21 cells (NEB, C2597). Cells were grown in Terrific Broth to mid-log phase, the temperature was reduced to 18 °C, and expression was induced at an OD₆₀₀ of 0.6 with 0.25 mM IPTG for 16–20 h before harvesting and freezing at −80 °C.

Cell pellets were resuspended in lysis buffer (50 mM HEPES, pH 7.5, 500 mM NaCl, 10% (v/v) glycerol, 1 mM TCEP) supplemented with a homemade protease inhibitor cocktail (1 mM PMSF, 5 µg/mL leupeptin, 1 µg/mL aprotinin, and 1 µg/mL pepstatin A). Cells were lysed using an LM20 microfluidizer (Microfluidics) or by sonication, and lysates were clarified by centrifugation at 34,000 × g for 30 min. Cleared lysates were incubated with Strep-Tactin Superflow Plus resin (Qiagen) at 4 °C for 1 h. The resin was extensively washed, and bound protein was eluted by on-resin SUMO cleavage with 10 µg home-made Ulp1 protease. Proteins were concentrated to 1 mg/mL, as measured using the Pierce 660 nm Protein Assay Reagent (Thermo Scientific) with a bovine serum albumin (BSA) standard curve, flash-frozen in liquid nitrogen, stored at −80 °C, and used for kinase assays.

Additional protein batches were further purified by ion-exchange chromatography. Cas13 and Csx34 was diluted to 100 mM NaCl, loaded onto an ÄKTA Pure system, and purified by heparin affinity chromatography (Heparin HP, Cytiva) using a linear gradient from 100 mM to 1 M NaCl. Peak fractions were collected, flash-frozen in liquid nitrogen, stored at −80 °C. Heparin purified Cas13 was used for tRNA cleavage sequencing, while heparin purified Csx34 was used in SELEX and fluorescence anisotropy assays. Both proteins were active in in vitro kinase assays.

### In vitro kinase reactions

Standard reactions (20 µL) were performed in 25 mM HEPES (pH 6.8), 5 mM MgCl₂, 0.05 µL murine RNase inhibitor (Thermo Scientific), 5% glycerol, 50 ng/µL salmon sperm DNA, 180 nM Cas13, 100 nM Csx33, 600 nM Csx34, and 2 mM ATP, supplemented with 0.25 µL [γ-³²P]ATP (3000 Ci/mmol, 10 mCi/mL; Revvity). The final NaCl concentration was 52 mM; RsCas13 cleavage efficiency decreased at NaCl concentrations above 60 mM. Reactions were incubated for 1 h at 37 °C and quenched by addition of 4× Laemmli sample buffer (Bio-Rad). Samples (10 µL) were resolved on 4–12% Bis-Tris SDS–PAGE gels (Novex, Invitrogen). Gels were fixed in 40% methanol/10% acetic acid for 1 h and washed for 4 h in water with frequent exchanges, then dried and exposed to a phosphor screen for 3 days (GE Healthcare). Phosphor screens were imaged using a Typhoon imager (GE Healthcare). Full-length tRNAs were added to reactions at a final concentration of 50 ng/µL, whereas small RNA fragments and CCA messengers were added at 10 ng/µL. Key experiments were performed independently at least twice, with similar results obtained. Representative results are shown unless otherwise indicated.

Components were omitted as indicated, and protein storage buffer was used to maintain a constant salt concentration. Lysates were treated for 30 min with proteinase K (37 °C, 15 min; NEB), DNase I (37 °C, 15 min; NEB), or immobilized RNase A bound to NHS-activated agarose (37 °C, 15 min; Pierce) to prevent RNAse contamination of the CASK reaction. Two-pot reactions were performed under identical buffer conditions, with each reaction incubated for 1 h. In the second step, 10 µL of the Cas13 reaction was combined with 10 µL containing Csx33, Csx34, and ATP.

### RNA-sequencing

A library of 47 *E. coli* full-length was cloned and purified as mentioned above. The tRNA library was pooled (2 µg total RNA) and subjected to Cas13 collateral digestion by mixing 1 µg Cas13 with 2 µg tRNA in reaction buffer containing 25 mM HEPES (pH 6.8), 50 mM NaCl, 5% (v/v) glycerol, 1 mM TCEP, and 5 mM MgCl₂, supplemented with equimolar crRNA and target RNA, and incubated at 37 °C for 1 h. Reactions were heat-inactivated at 85 °C for 5 min and treated sequentially with 1 µL RppH (37 °C, 30 min) and 1 µL T4 PNK (37 °C, 6 h), followed by addition of 1 µL T4 PNK with 10 mM ATP and incubation for 1 h at 37 °C further. RNA was purified using RNAClean XP beads and eluted in 10 µL. Small RNA-seq libraries were prepared using the NEBNext Small RNA Library Prep Set for Illumina (NEB) according to the manufacturer’s instructions and sequenced on an Illumina NextSeq 2000 to obtain approximately 5 million reads per sample. Reads were trimmed to remove kit-derived 5′ and 3′ flanking sequences, and inserts were aligned to a synthetic *E. coli* tRNA reference using srnaMapper (35). 3′-end positions were counted and aggregated, and the z-score of the 3′-end enrichment was calculated using a custom Python script.

### LC-MS analysis

CCA fragments were analyzed by LC–MS using 5 µL injections separated on a Waters BEH Z-HILIC column (2.1 × 100 mm) with a flow rate of 0.3 mL/min at 40 °C. Mobile phase A consisted of 10 mM ammonium formate and 0.1% (v/v) formic acid in water, mobile phase B consisted of 10 mM ammonium formate and 0.1% (v/v) formic acid in acetonitrile. A gradient from 99% to 30% B was run over 10 min, followed by wash and re-equilibration. Analytes were detected by electrospray ionization in full-scan positive-ion mode on a Thermo Scientific Vanquish HPLC coupled to an Exploris Orbitrap mass spectrometer. Positive control samples were prepared from a synthetic RNA oligonucleotide (GGGCCA) digested with RNase T1. Reaction samples were prepared in 25 mM HEPES (pH 6.8), 50 mM NaCl, 1 mM TCEP, and 5 mM MgCl2 and assembled as RNP complexes using 1 µg protein with equimolar crRNA and target RNA, with 2 µg tRNA^Gly(CCC)^ included in reactions with or without target RNA. For CCA binding assays, purified protein was immobilized on Ni-NTA resin, incubated with RNase T1-generated CCA fragments, washed, eluted with imidazole, and eluates were analyzed by LC–MS under the conditions described above.

### Csx34 SELEX

Random DNA libraries were synthesized with a 24-nt randomized region flanked by next-generation sequencing handles. The initial library was amplified using Taq DNA polymerase (NEB) according to the manufacturer’s protocol with 12 PCR cycles. PCR products were evaluated by agarose gel electrophoresis to confirm a single species without detectable heteroduplexes. The amplified library was purified using a DNA Clean & Concentrator-25 column (Zymo Research) and used for downstream SELEX experiments.

Experiments were performed using SELEX buffer containing 25 mM HEPES (pH 7.5), 50 mM NaCl, 1 mM MgCl₂, and 4% (v/v) glycerol. Binding reactions were carried out in SELEX buffer supplemented with 1 mM TCEP and 5 µg/mL poly(dI-dC), and wash steps were performed in SELEX buffer supplemented with 1 mM TCEP. For round 1, 1 µg of His-tagged target protein was incubated with 1.5 µg of the SELEX DNA library and captured using Ni-NTA magnetic beads (Thermo Fisher) for 30 min at room temperature, followed by three washes. To recover enriched sequences, 1 µL of bead slurry was used as input for PCR amplification with Taq DNA polymerase (12 cycles for round 1 and 14 cycles for all subsequent rounds). After each round, PCR products were purified using a DNA Clean & Concentrator-5 column (Zymo Research); 200 ng of purified DNA was carried forward into the next round, and 1 µL of each purified library was retained for sequencing library construction.

Sequencing libraries were generated by PCR amplification of each round’s retained library using customized Illumina-compatible barcoded primers. For each round, input DNA was normalized, and 100 ng of library was used for amplification. Each reaction was performed with two replicates, starting with independent DNA libraries. Final libraries were sequenced on an Illumina MiSeq. Sequencing data were analyzed by trimming adapter and flanking sequences. The enriched 24-bp libraries were searched against the negative control using MEME with discriminative mode and default parameters^42^.

### Fluorescence anisotropy

Fluorescence anisotropy (fluorescence polarization) assays were performed using 15-nt oligonucleotides bearing a 5′ fluorescein label (IDT). Labeled oligonucleotides were annealed to reverse-complement strands in IDT Duplex Buffer. Binding reactions were assembled in 25 mM HEPES (pH 7.5), 50 mM NaCl, 5 mM MgCl2, 1 mM TCEP, and 5 µg/mL poly(dI-dC), using 5 nM duplex substrate and titrated protein (prepared at 500 nM and serially diluted). Reactions (80 µL total volume; 60 µL protein dilution + 20 µL substrate) were incubated at 37 °C for 30 min and read on SpectraMax iD5e Multi-Mode Microplate Reader (Molecular Device) in fluorescence polarization mode with G = 1, in FAM channel. Signals were background-corrected by subtracting the polarization of labeled substrate alone. Each concentration was measured in three technical replicates, and curves were plotted in GraphPad Prism. Key experiments were performed independently with separate protein batches, with similar results obtained.

### Mass photometry

Mass photometry samples were prepared using a 300-bp DNA substrate generated by PCR (with or without the identified Csx34 motif) and purified using AMPure XP beads (Beckman Coulter). Wild-type Csx34 or Csx34 T269E protein was incubated with DNA in buffer containing 25 mM HEPES (pH 7.5), 50 mM NaCl, 5 mM MgCl2, 1 mM TCEP, and 5 µg/mL poly(dI-dC) at 37 °C for 30 min prior to measurement. Samples were diluted to 10 nM on coverslip and measured by droplet dilution mode on Refeyn TwoMP mass photometer. Movies were acquired for 1 min (∼1,000 events) and analyzed using Refeyn MP Discovery software to generate mass histograms. Each condition was measured at least twice, with similar results obtained.

### Cryo-EM sample preparation and data collection

Csx34 T269E was purified as described above, except that glycerol was omitted during the Heparin HP step and all procedures were performed in cryo-EM buffer (25 mM HEPES pH 7.5, 150 mM NaCl, 2 mM MgCl₂). Two 15-nt motif-containing reverse-complementary oligos were synthesized by Azenta and annealed in DNase-free water at 50 μM by heating to 75 °C for 5 minutes and slowly cooling to 4 °C. Heparin-purified Csx34 T269E (2 mg mL⁻¹) was incubated with the 15-bp motif-containing dsDNA at a protein:DNA ratio of 1:1.2 for 30 min at 25 °C and further purified through a Superose 6 column (Cytiva). Peak fractions were harvested and concentrated. Immediately before application to Quantifoil R1.2/1.3 300-mesh Au holey-carbon grids (Quantifoil), 3.5 μL of the Csx34 T269E–DNA complex at 1 mg/mL was mixed with 1 μM fluorinated Fos-Choline-8 (Anatrace). Grids were then blotted and plunge-frozen immediately in liquid ethane using a Vitrobot Mark IV (Thermo Fisher Scientific).

Data were collected on a Titan Krios microscope operated at 300 kV and equipped with either a Gatan K3 direct electron detector or a Falcon 4 detector (Thermo Fisher Scientific). Movies were recorded using Legion software at a physical pixel size of 0.649 Å per pixel, with 5-s exposures fractionated into 50 frames and a total dose of ∼57.04 e⁻ Å⁻²

### Image processing and 3D reconstruction

Dark-subtracted movies were gain normalized. Motion correction and dose weighting were performed in Legion using MotionCor2. The contrast transfer function (CTF) was estimated with PatchCTF in cryoSPARC^43^. Particles were picked using a customized trained cryolo model, and a total of 1,793,974 particles were imported into cryoSPARC. After removing duplicates within 100 Å, a total of 1,096,678 particles were extracted in cryoSPARC with a box size of 512 pixels and downsampled to 256 pixels to accelerate processing. Given that the size of the Csx34 T269E– DNA complex is less than 80 kDa, we adapted the HR-FAIR method to align extracted particles using only high-resolution signal during 2D classification and Ab-initio refinement.

We customized cryoSPARC 2D classification with the following settings: 200 2D classes, maximum resolution 3 Å, maximum alignment resolution 3 Å, initial classification uncertainty factor 1, circular mask diameter 70 Å, minimum separation distance 100 Å, number of final full iterations 20, and number of online-EM iterations 80. After 12 hours of 2D classification, the results converged around 12 full iterations, and the classification was manually stopped. Classes showing clear orientations of single particles were selected, yielding a total of 291,060 particles.

The selected particles were subjected to Ab-initio reconstruction with the following customized settings: number of Ab-initio classes 3, maximum resolution 2.6 Å, initial resolution 5 Å, Fourier radius step 0.005, center structures in real space OFF, initial minibatch size 300, and final minibatch size 1000. Ab-initio refinement lasted more than 28 hours, and among the three classes, the one with clearer secondary structure was selected, yielding a total of 73,024 particles.

The selected particles were re-extracted from the micrographs with a 512-pixel box size and reconstructed without changing the alignment information using the Reconstruct Only function in cryoSPARC. 3D refinement was further carried out with one round of local refinement using the following customized settings: rotation search extent 2 degrees, shift search extent 1 degree, and initial low-pass resolution 6 Å. This processing yielded a final map at 3.12 Å resolution with clear secondary structural features of the protein and DNA.

### Cryo-EM model building and refinement

The refined map was further post-processed using EMReady2 to facilitate model building. The initial model was built using the unprocessed map with ModelAngelo^44^, and the correct regions from the raw output were selected. The post-processed map was then used for further atomic model building manually in Coot^45^, given that none of the structural prediction models could faithfully predict the Csx34 T269E–DNA complex. Ambiguous side chains were pruned, and several solvent-contacted loops were not built because of insufficient density. The final model was refined in real space using Phenix^46^. For cross-validation, the final model was refined against one half-map, and model–map Fourier shell correlation (FSC) curves were generated using the comprehensive validation module in Phenix. Phenix and MolProbity^47^ were used to validate the final model. Structure figures were generated in ChimeraX.

### Bacterial reporter, phage defense, and RNA sensing assays

Luciferase reporter plasmids containing the native *Ruminococcus* upstream regulatory sequence or the indicated mutagenized variants were co-transformed into *E. coli* 5α cells (NEB, C2987) with Csx33/Csx34 helper plasmid variants or an empty vector. Individual colonies were inoculated as independent biological replicates. Overnight cultures were diluted to an OD₆₀₀ of 0.05 and incubated at 37 °C with shaking for 4 h before luminescence was measured using a SpectraMax iD3e Multi-Mode Microplate Reader (Molecular Devices).

For synthetic transcriptional assays with Csx34–SoxS fusion constructs, luciferase reporter plasmids containing the Csx34 SELEX motif or a randomized control sequence upstream of the J23112 promoter were introduced into *E. coli* BL21 cells (NEB, C2597) together with plasmids encoding Csx34–SoxS fusion variants or an empty vector. Cultures were prepared as described above, except that after 2 h of growth, aTc was added to a final concentration of 10 ng/mL and incubation continued for an additional 3 h before luminescence measurement.

For T7 phage experiments, we generated a reporter plasmid containing the native *Ruminococcus* regulatory sequence upstream of LbuCas13. Reporter plasmids were co-transformed into *E. coli* strain C cells (ATCC) with Csx34 helper plasmid variants or an empty vector. Overnight cultures were diluted 1:20 in 0.75% top agar, and 5-fold dilutions of T7 phage (ATCC) spotted.

For in vivo RNA-sensing assays, *E. coli* 5α cells (NEB, C2987) were co-transformed with a plasmid encoding RsCas13 and either a targeting or non-targeting guide RNA, together with a luciferase reporter plasmid containing a promoter mutation (P1) in the *Ruminococcus* upstream regulatory sequence, *luxCDABE*, and a target RNA transcript complementary to the guide RNA. Transformed cells were made competent using the Mix & Go! Competent Cell Kit (Zymo Research), and 50 µL of competent cells were transformed with Csx33/Csx34 helper plasmid variants or an empty vector control. Individual colonies were used as independent biological replicates. Overnight cultures were diluted to an OD₆₀₀ of 0.05 and incubated at 37 °C with shaking for 2.5 h before luminescence was measured.

## Acknowledgements

We thank Ross Tomaino and the Harvard Medical School (HMS) Taplin Mass Spectrometry Facility for phosphoproteomic analysis, and Michael James and the HMS Analytical Chemistry Core for LC–MS analysis. Structural work was performed at the National Center for Cryo-EM Access and Training (NCCAT) and the Simons Electron Microscopy Center located at the New York Structural Biology Center with the assistance from Zephan Melville, Mahira Aragon, Collin McManus, supported by National Institutes of Health (Common Fund U24GM129539, NIGMS R24GM154192), the Simons Foundation (SF349247), and the NY State Assembly. J. S. is supported by the National Institutes of Health Director’s New Innovator Award (DP2-HL185100).

## Author contributions

J.S., Y.L., Y.Q., I.B., A.Y., and H.C. performed experiments. J.S., Y.L., and Y.Q. wrote the manuscript. J.S. supervised the research.

## Data availability statement

Data is available upon request. Plasmids are available from Addgene. RNA sequencing reads and SELEX library sequencing reads are available at the NCBI Sequence Read Archive with Project ID PRJNA1422099. The Csx34 T279E–DNA cryo-EM map has been deposited in the Electron Microscopy Data Bank with code EMD-76409. The coordinates for the composite atomic model have been deposited in the Protein Data Bank under accession code 12FR. Further inquiries and material requests should be directed to the lead contact, Jonathan Strecker.

## Code availability statement

Scripts used to analyze the RNA-seq data and generate the heat map have been deposited at Zenodo with 10.5281/zenodo.18599311.

---

**Supplementary Table 1.** List of CASK loci and proteins

**Supplementary Table 2.** Proteins used in this study

**Supplementary Table 3.** Nucleic acids sequences used in this study

**Supplementary Table 4.** Cryo-EM data collection, refinement, and validation statistics

**Extended Data Figure 1.**
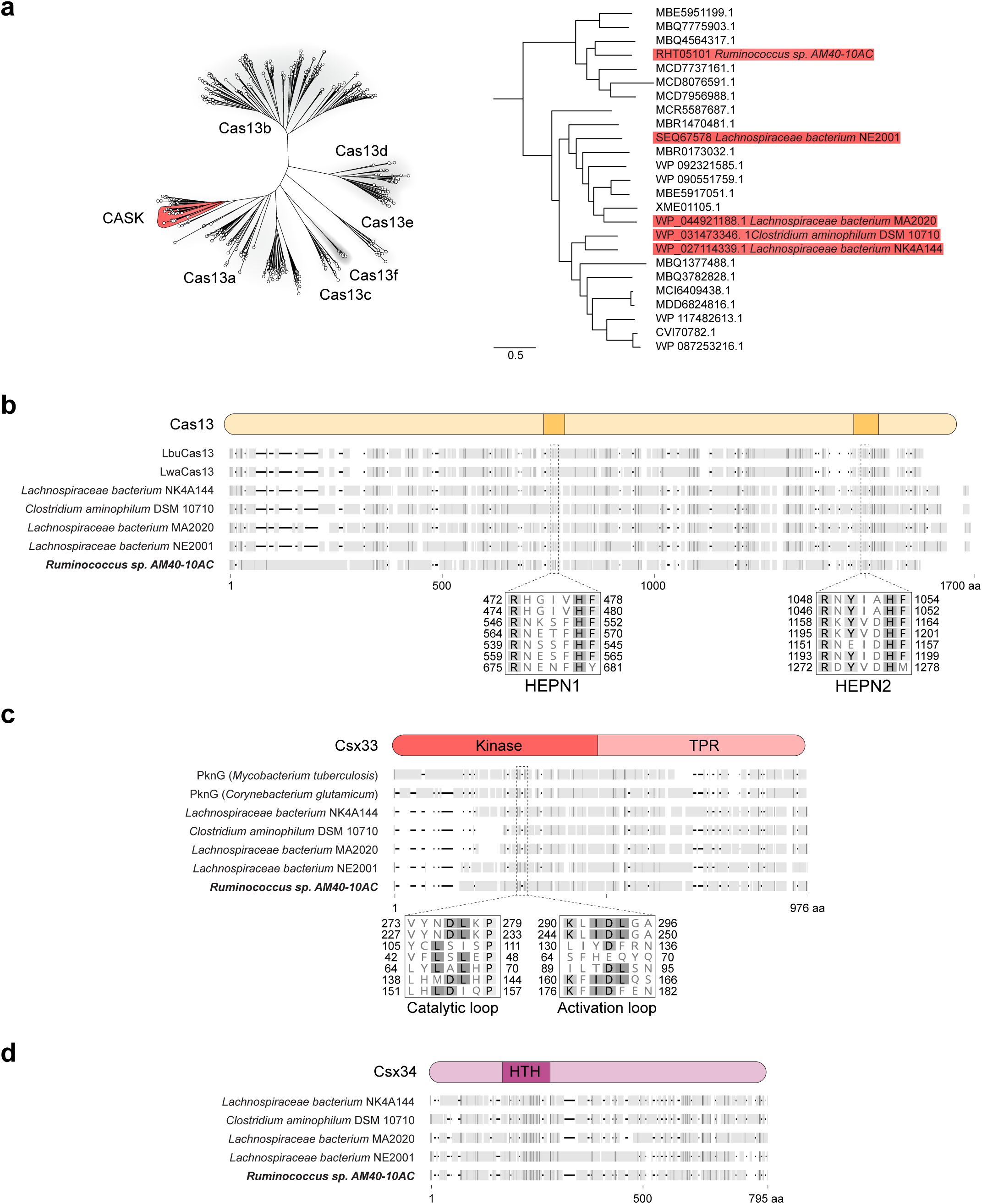
Analysis of CASK protein components. **a**, Tree of Cas13a proteins. Cas13a proteins encoded within CASK systems are indicated in red. **b,** Alignment of Cas13 nucleases with highlighted HEPN domains. **c,** Alignment of Csx33 kinases with highlighted catalytic and activation loops. **d,** Alignment of Csx34 proteins.

**Extended Data Figure 2.**
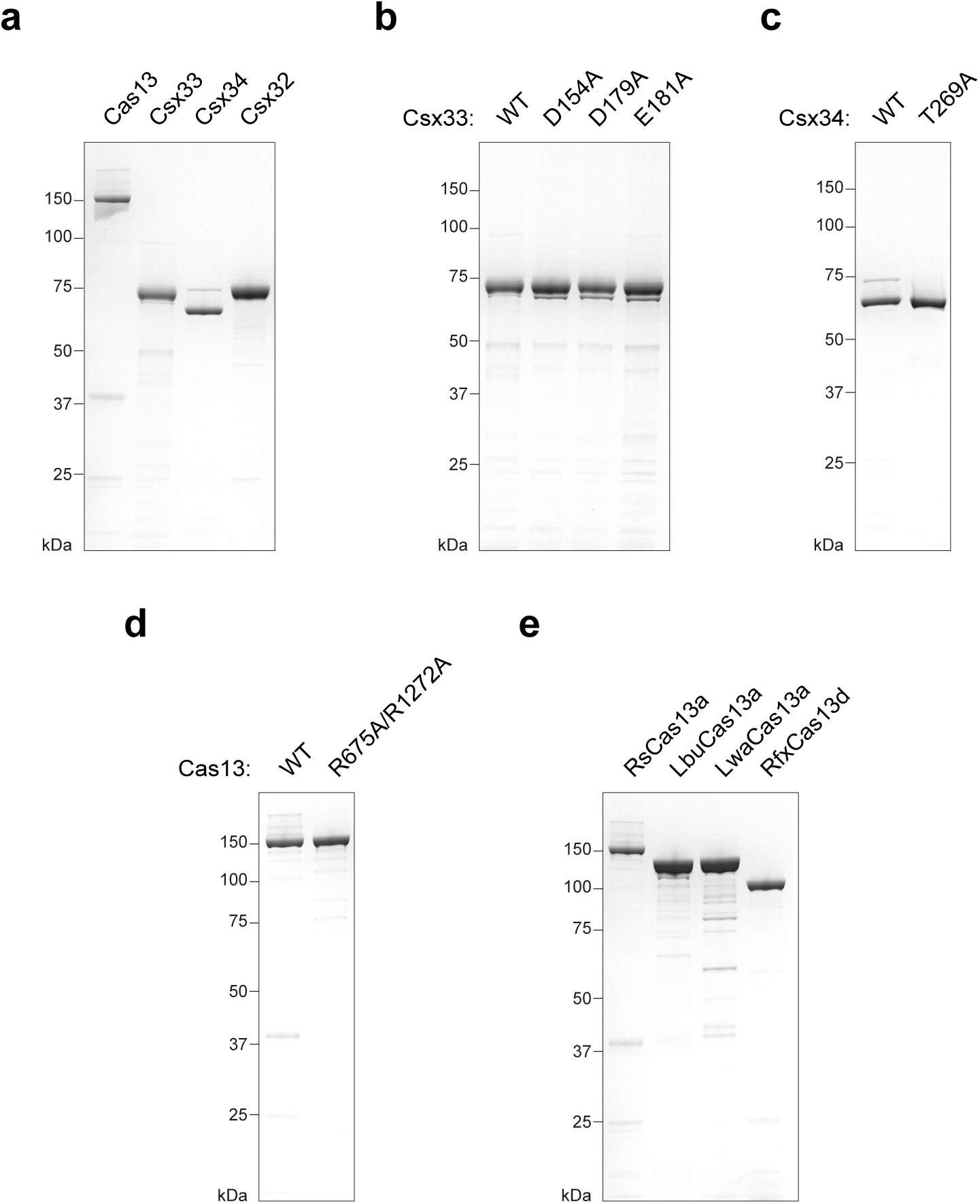
Proteins used in this study. **a**, RsCASK proteins from *Ruminococcus sp. AM40-10AC*. **b,** Csx33 kinase mutants. **c,** Csx34 phosphorylation site mutant. **d,** RsCas13 HEPN catalytic mutant. **e,** Cas13 orthologs. (a–e) are SDS–PAGE gels stained with Coomassie blue.

**Extended Data Figure 3.**
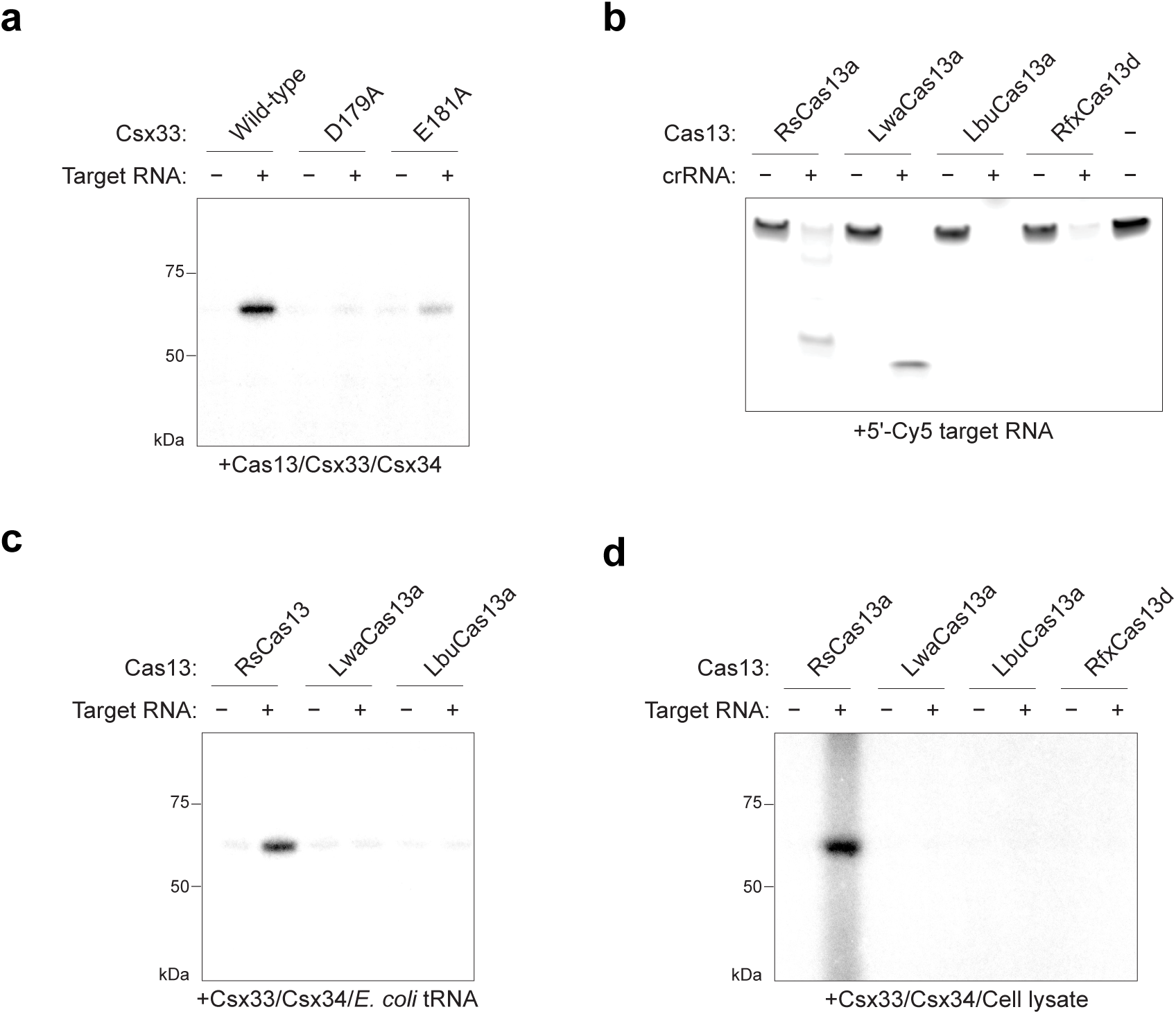
CASK in vitro reactions. **a**, Effect of Csx33 mutations on CASK activity. **b,** Target cleavage of a Cy5-labeled RNA by Cas13 orthologs used in this study. **c, d,** Ability of diverse Cas13 orthologs to activate Csx33 in reactions with *E. coli* tRNA (c) and *E. coli* cell lysate (d).

**Extended Data Figure 4.**
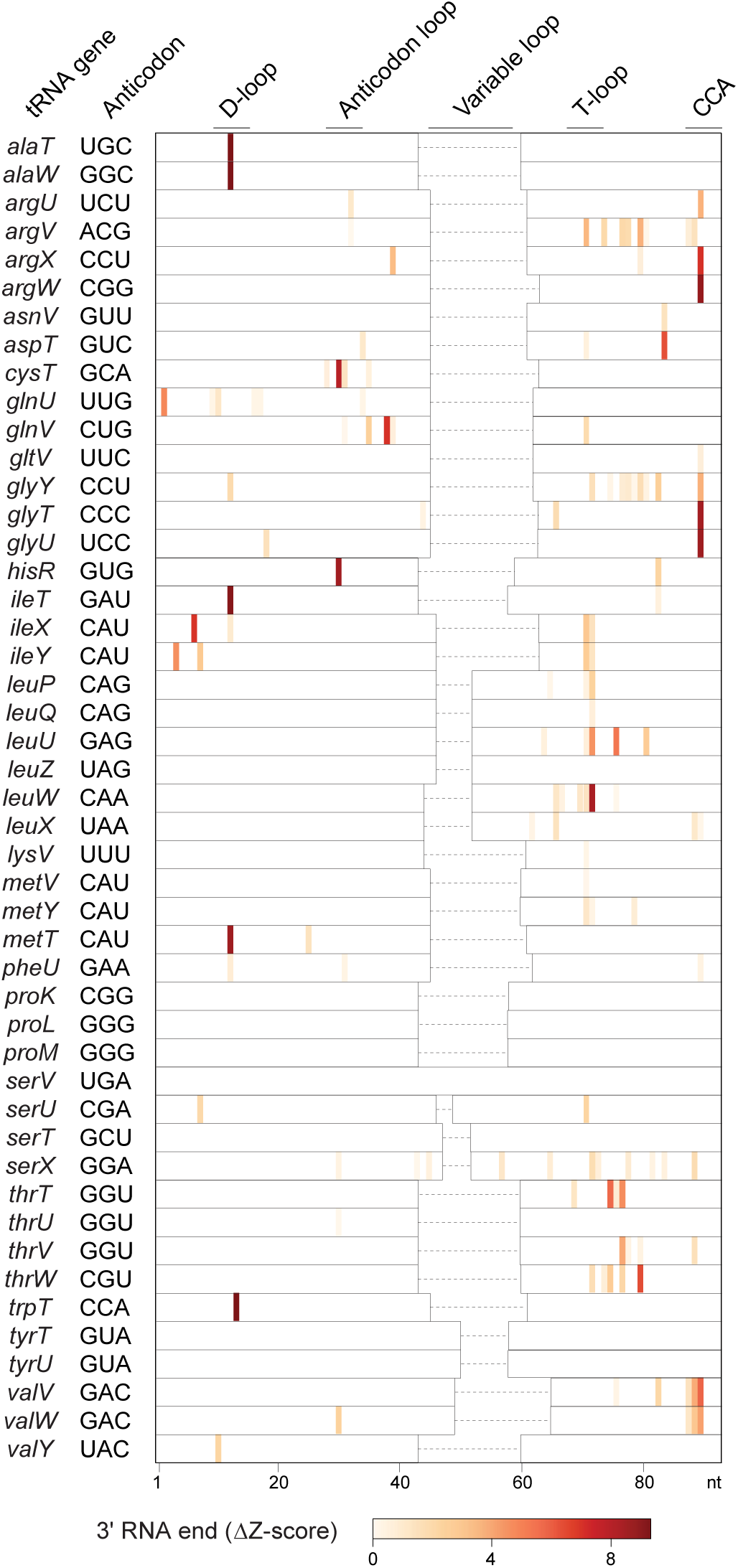
RsCas13 collateral RNA sequencing. RsCas13 collateral cleavage of in vitro–transcribed *E. coli* tRNAs. Novel 3′ tRNA ends detected by RNA sequencing following in vitro reactions containing target RNA, compared to reactions with random RNA as a control. (a) and (c) are autoradiographs of SDS–PAGE-resolved in vitro kinase reactions.

**Extended Data Figure 5.**
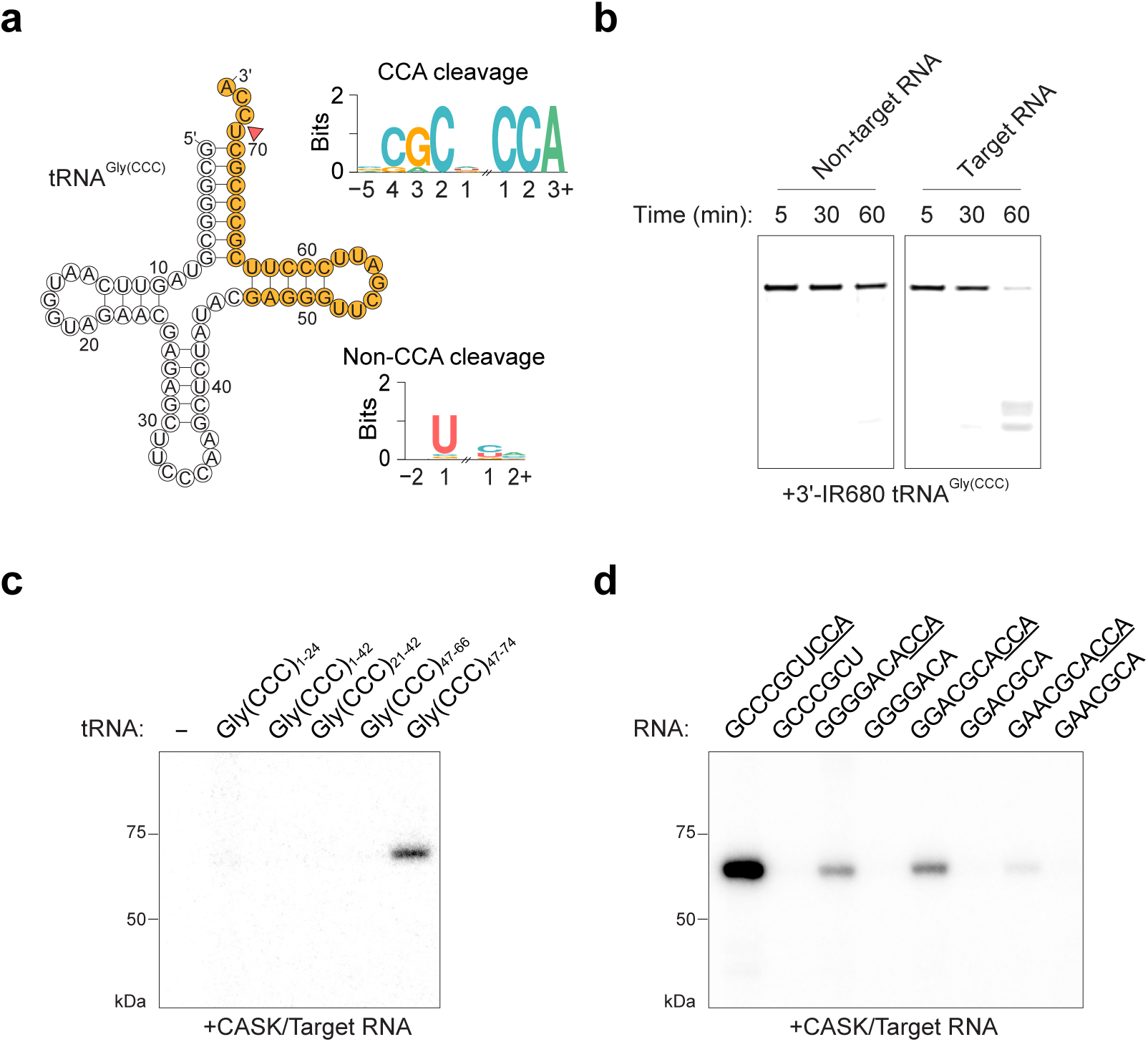
RsCas13 cleaves tRNA to activate Csx33. **a**, Schematic of tRNA^Gly(CCC)^ cleaved by RsCas13 (red arrow) to generate CCA trinucleotides and nucleotide motif analysis of cleaved tRNAs identified by RNA sequencing. A truncated tRNA sequence that is able to support CASK activity in (c) is highlighted in yellow. **b,** In vitro collateral cleavage of tRNA^Gly(CCC)^ by RsCas13 in response to target RNA. **c,** Mapping of a sequence in tRNA^Gly(CCC)^ that is sufficient for CASK activity, highlighted in (a). **d,** CASK activity is dependent on the presence of 3′ CCA sequence at the end of tRNA fragments (underlined). (c) and (d) are autoradiographs of SDS–PAGE-resolved in vitro kinase reactions.

**Extended Data Figure 6.**
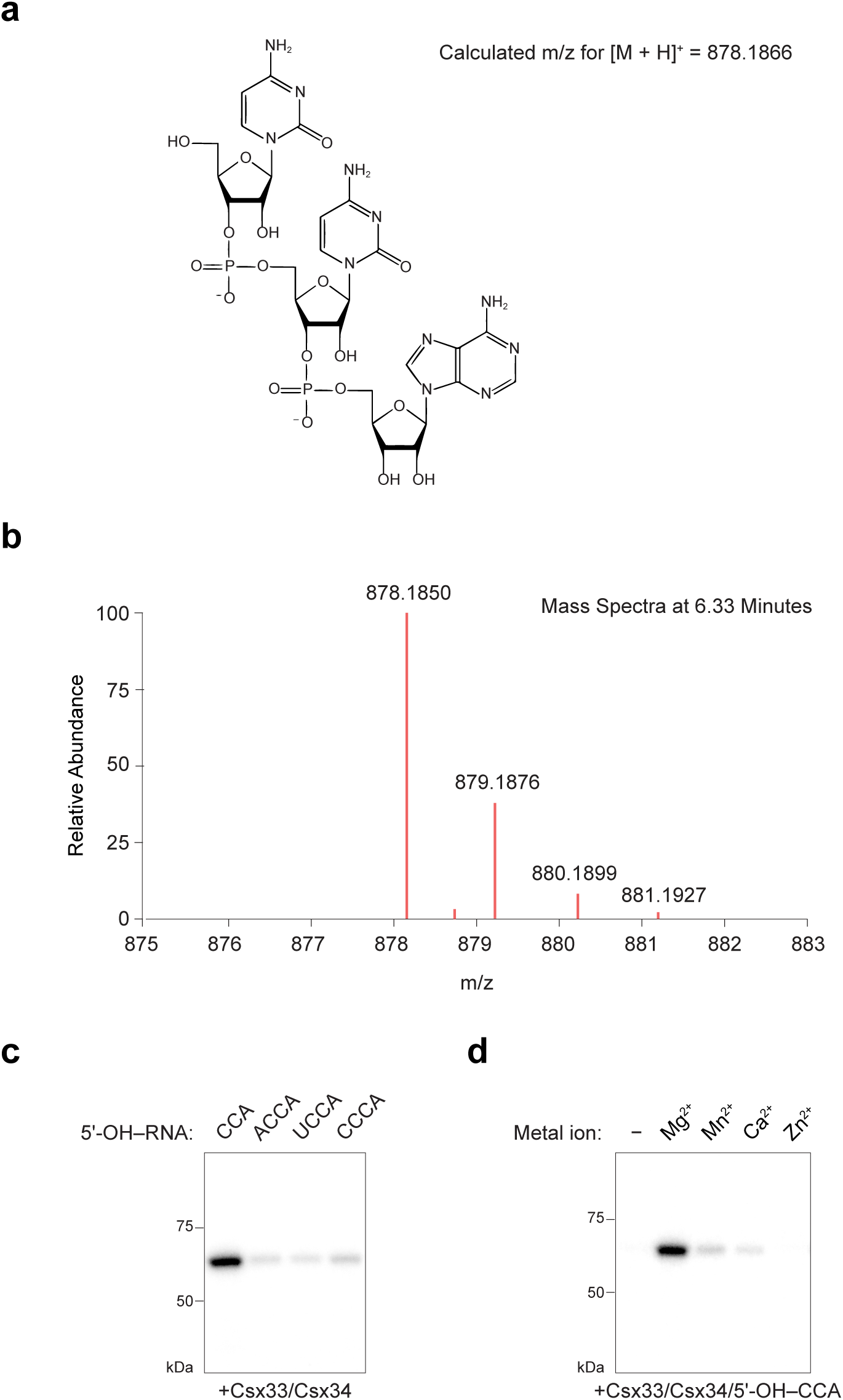
Liquid chromatography–mass spectrometry analysis of CCA trinucleotides. **a**, Chemical structure and predicted mass-to-charge ratio (m/z) of 5′-OH–CCA trinucleotides. ESI–MS: calculated *m/z* for [M + H]+ (C_28_H_37_N_11_O_18_P_2_) = 878.1866. **b,** MS spectrum of the peak at 6.33 min with an observed *m/z* of 878.1850. **c,** Csx33 activity with tetranucleotides generated by RNase T1 digestion. **d,** Csx33 activity with 5′-OH–CCA and different divalent metal ions. (c) and (d) are autoradiographs of SDS– PAGE-resolved in vitro kinase reactions.

**Extended Data Figure 7.**
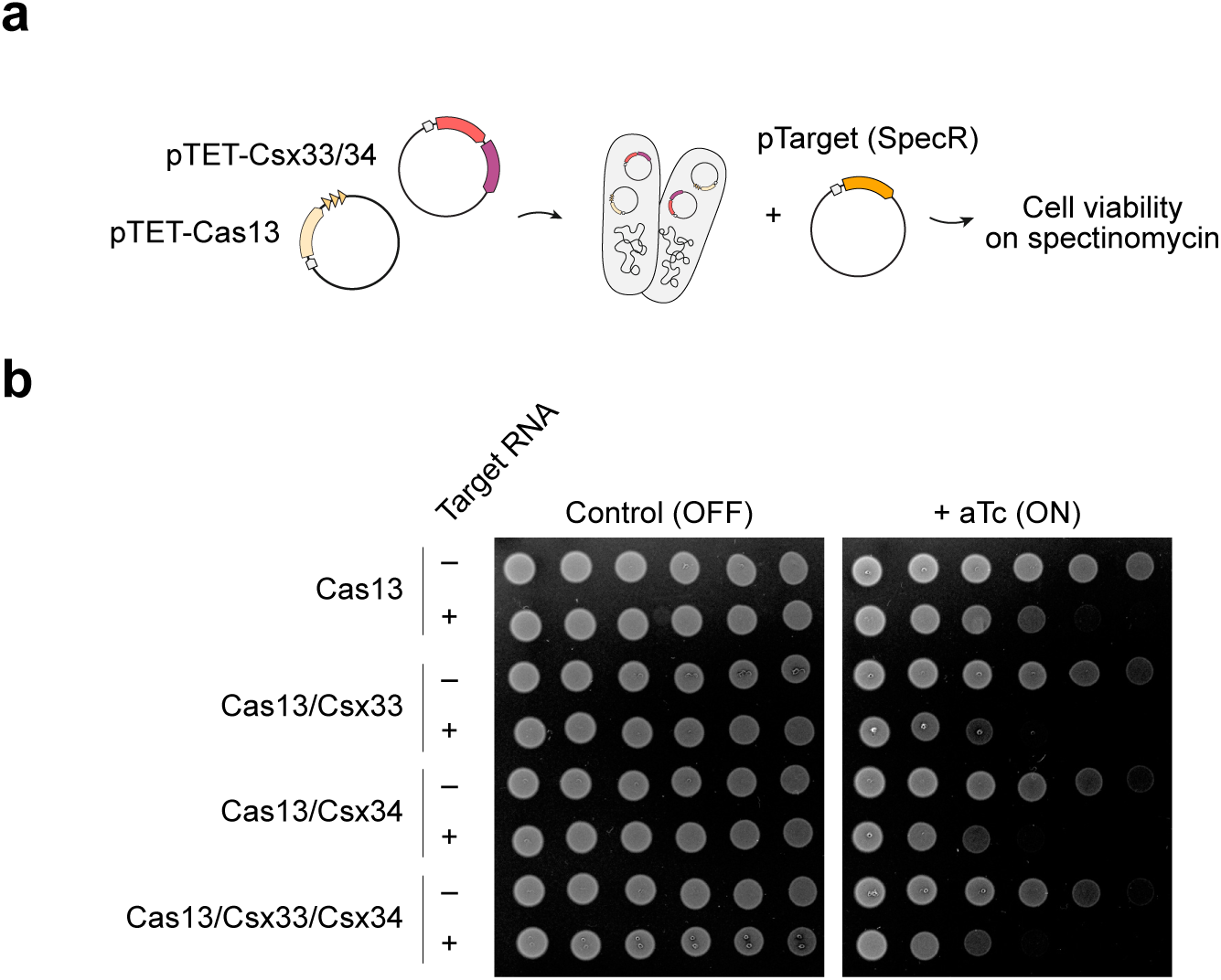
Impact of CASK genes on RsCas13-mediated defense. **a**, Schematic of an interference assay in *E. coli* expressing RsCas13 and Csx33–Csx34. Cells were transformed with a target plasmid (pTarget) encoding a spectinomycin resistance RNA that matches the Cas13 crRNA, or a variant containing synonymous substitutions in the target that escapes recognition. **b,** RsCas13 provides interference against a spectinomycin resistance transcript, while co-expression of Csx33 and Csx34 shows no substantial effect on interference. Cells were serially diluted twofold and spotted on spectinomycin plates without aTc (OFF) or on plates containing 1 ng/mL aTc (ON) to induce CASK protein expression.

**Extended Data Figure 8.**
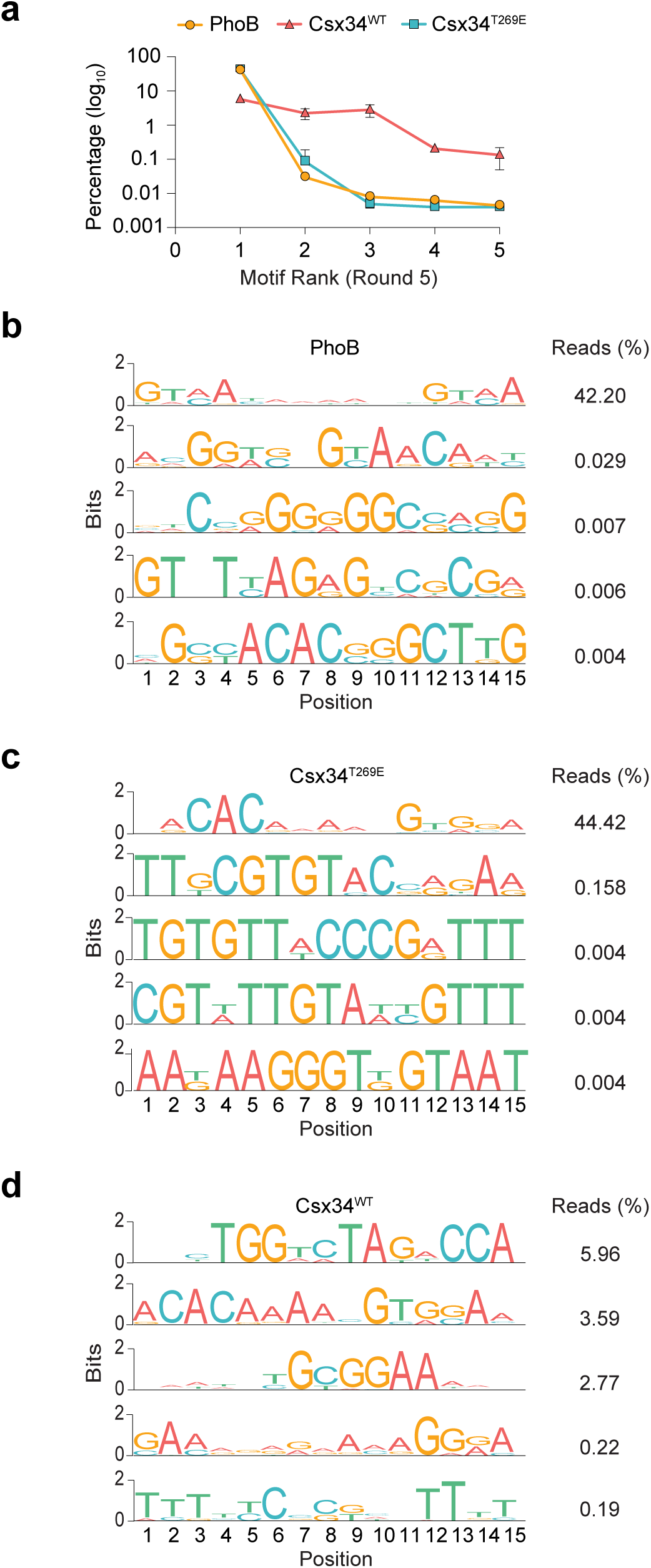
Identification of a Csx34 binding motif by SELEX. **a**, Identification of a single highly enriched motif after five rounds of SELEX for Csx34 T269E, but not wild-type Csx34. PhoB was used as a positive control. **b–d,** The top five motifs identified after five rounds of SELEX and the corresponding number of reads, shown as a percentage of the total, for the PhoB positive control (b), Csx34 T269E (c), and wild-type Csx34 (d).

**Extended Data Figure 9.**
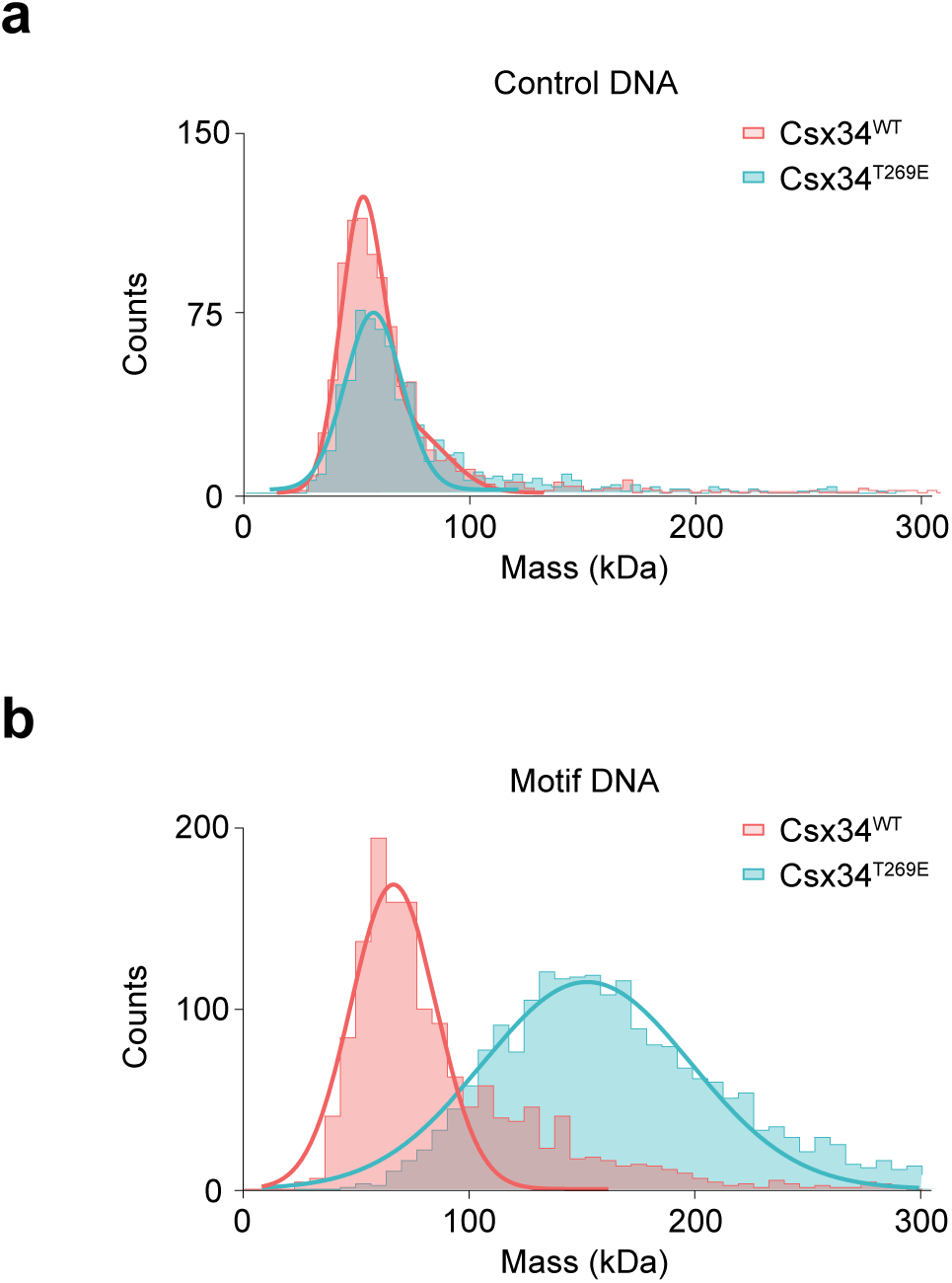
DNA binding of Csx34 measured by mass photometry. **a, b**, Binding of wild-type Csx34 and Csx34 T269E protein to a control DNA substrate (a) or to a substrate containing the identified motif (b).

**Extended Data Figure 10.**
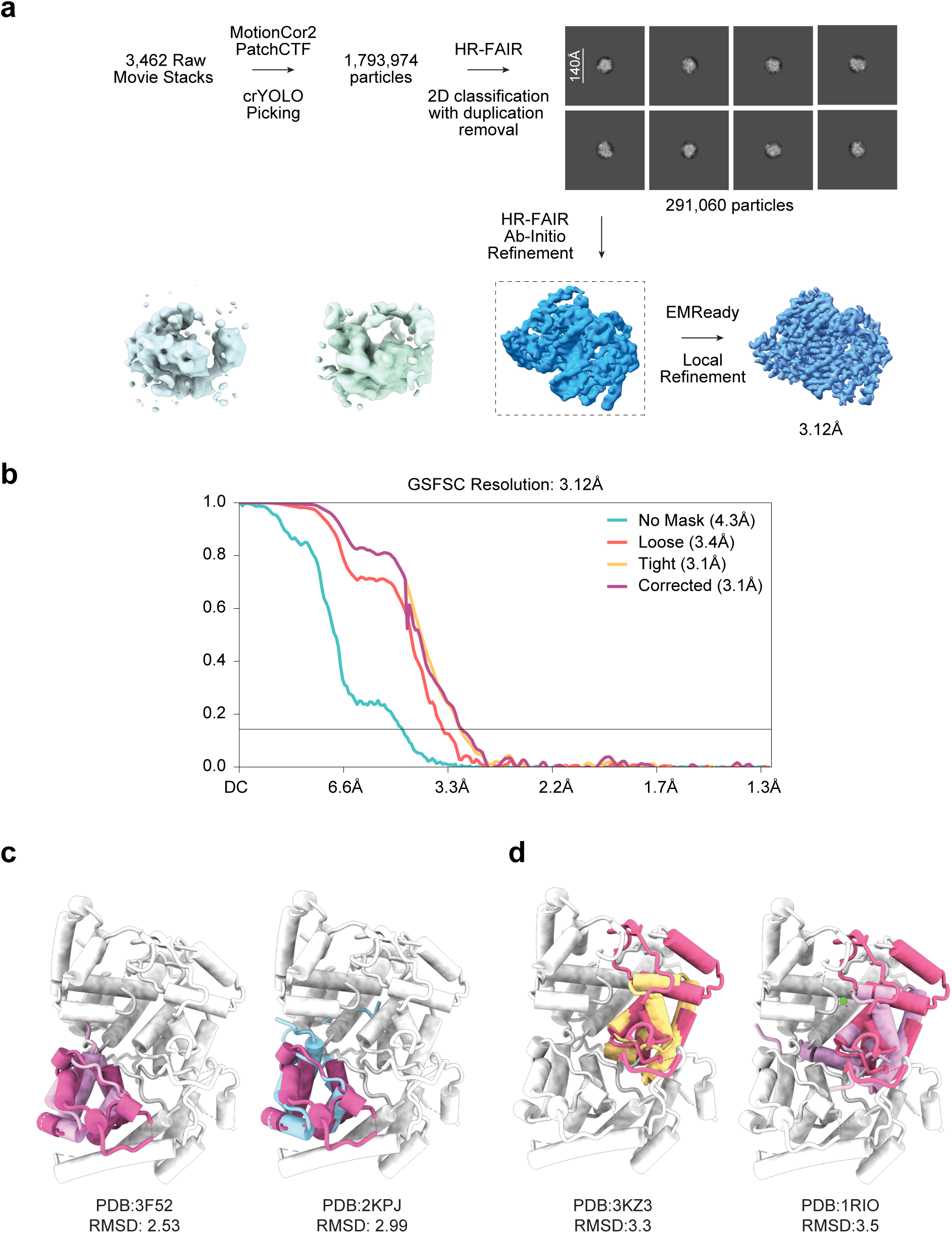
Data processing of a Csx34 T269E–DNA complex. **a**, Data-processing workflow. Maps generated from HR-FAIR Ab-Initio refinement are shown. The final CryoSPARC local-refinement map had an overall resolution of 3.12 Å based on the half-map FSC. The EMReady post-processed map was used for manual model inspection/visualization. **b,** Fourier shell correlation curve calculated from the CryoSPARC local-refinement half-maps. **c, d,** Structural searches and alignment of HTH1 (c) and HTH2 (d) to characterized HTH domains.

**Extended Data Figure 11.**
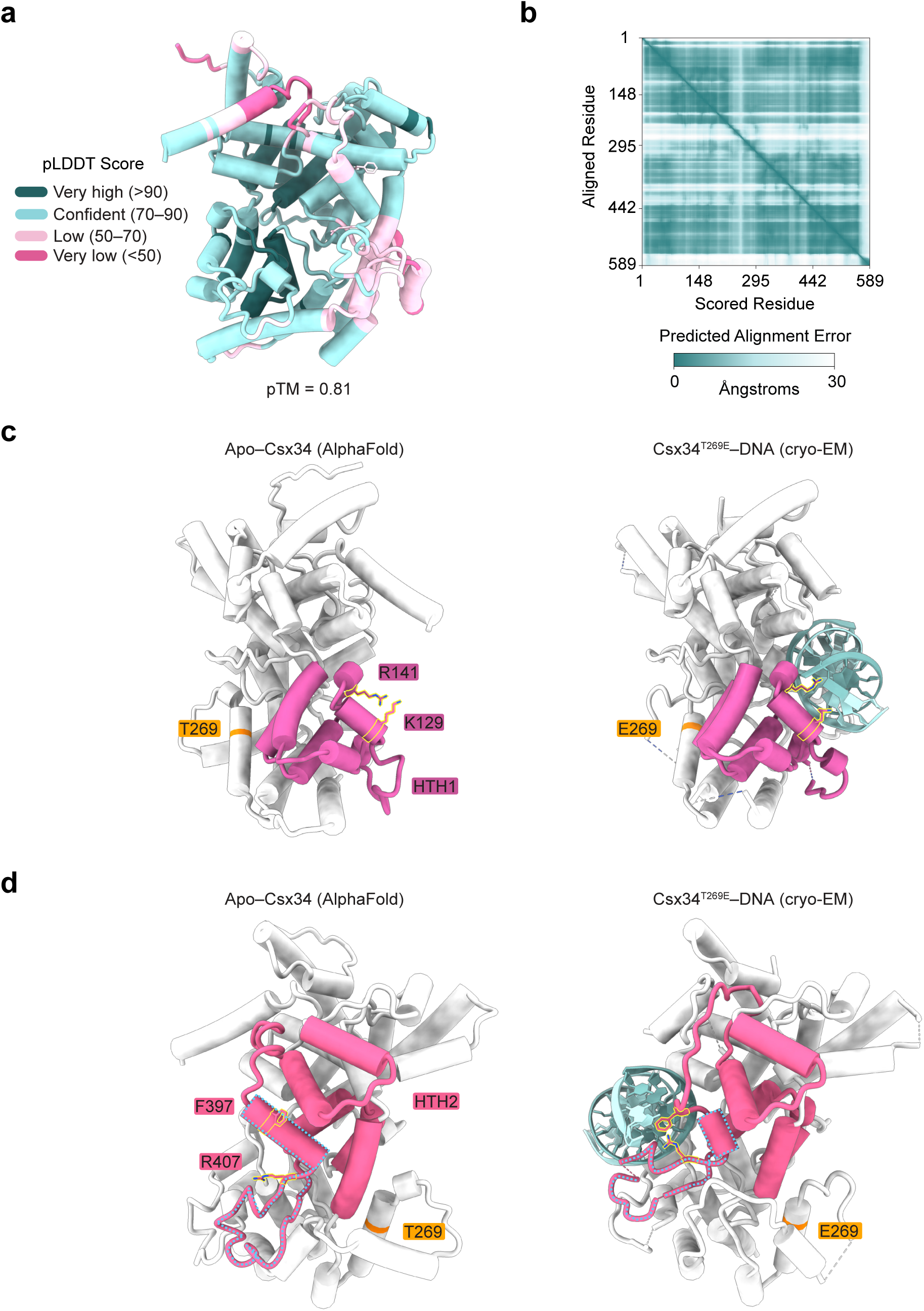
Predicted structure of apo–Csx34. **a**, AlphaFold 3 model of apo–Csx34 and the pLDDT score per position. **b,** Predicted Aligned Error (PAE) plot of the apo–Csx34 model. **c,** Comparison of HTH1 between the predicted apo-Csx34 model (c) and the experimental structure of Csx34 T269E–DNA (d), reveals no major conformational changes. **d,** Comparison of HTH2 between the predicted apo-Csx34 model (e) and the experimental structure of Csx34 T269E–DNA (f) revealing rearrangement of HTH2 and the repositioning of Phe397 and Arg407.

**Extended Data Figure 12.**
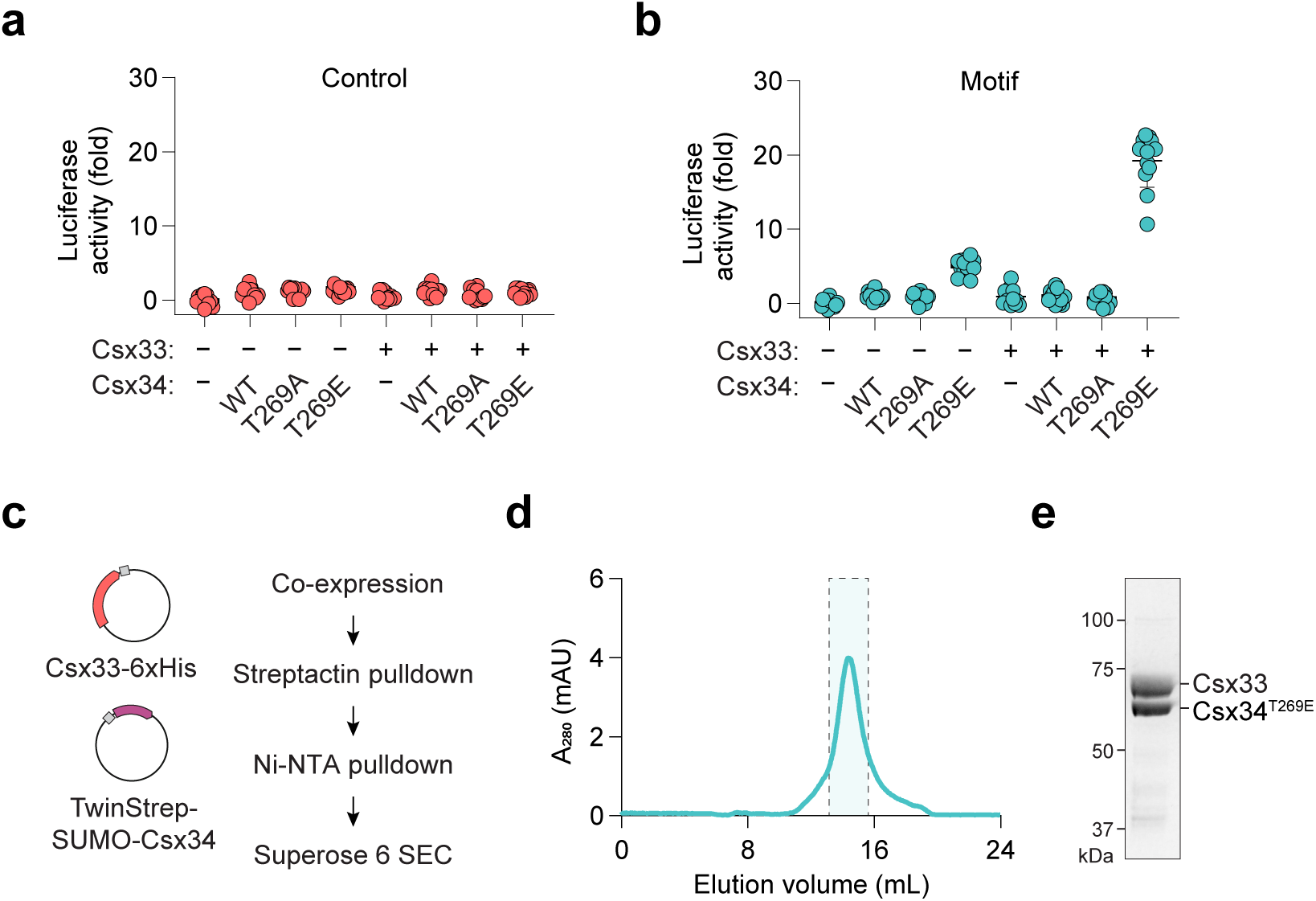
Transcriptional activation by a Csx33–Csx34 complex in *E. coli*. **a, b**, Luciferase activity following expression of Csx33 and Csx34 and a reporter containing the native *Ruminococcus sp.* sequence (b) or a control in which the Csx34 binding motif was mutated (a). Expression was normalized to cells carrying an empty control plasmid; error bars represent the standard deviation from the mean. *n* = 12 replicates. **c,** Schematic of co-purification strategy to isolate a Csx33–Csx34 complex from *E. coli*. **d,** Chromatogram of a Csx33–Csx34 T269E complex run on a Superose 6 Increase gel filtration column. **e,** SDS-PAGE analysis of a co-eluting Csx33–Csx34 T269E complex from (d).

**Extended Data Figure 13.**
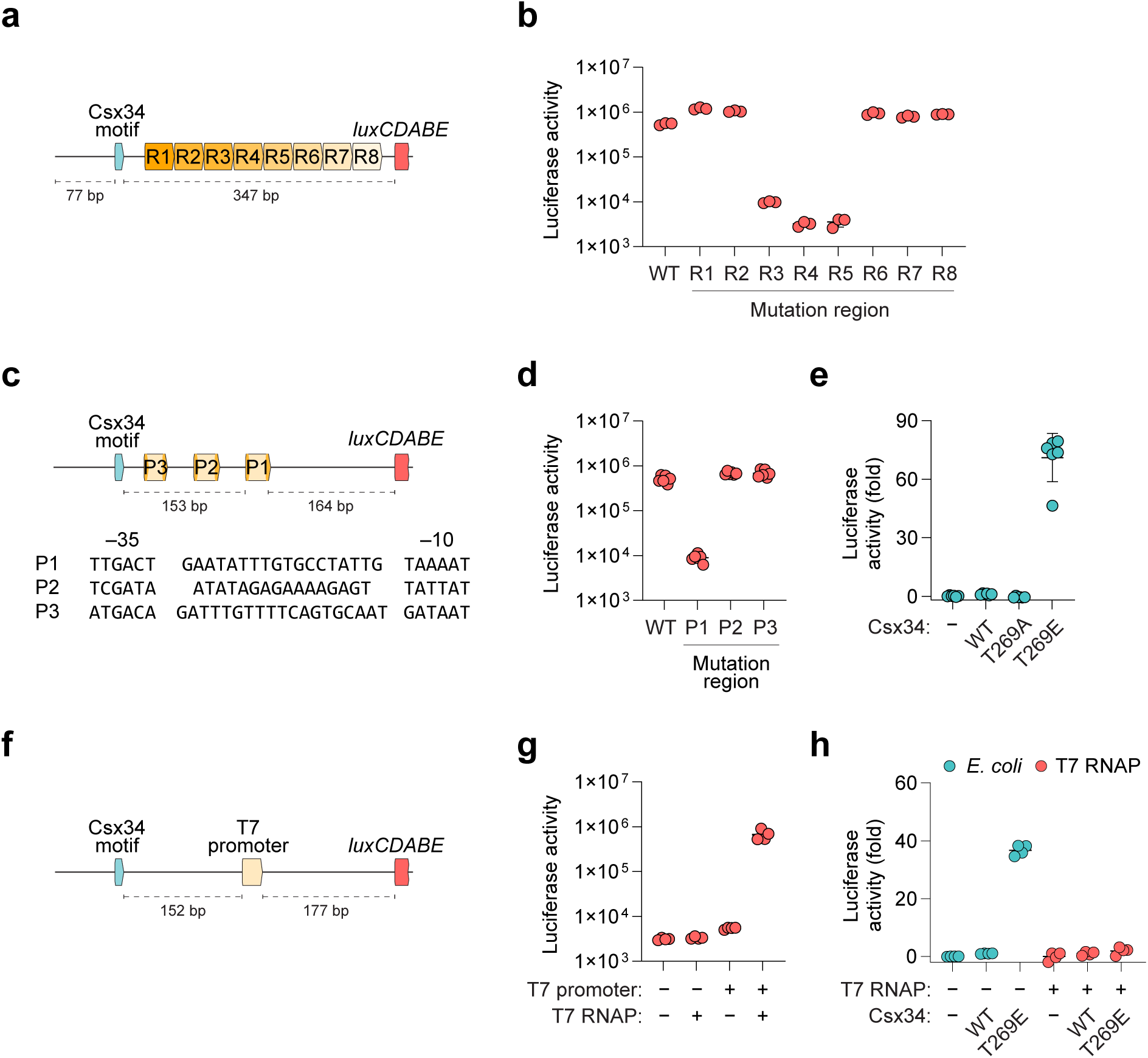
Investigation of the *Ruminococcus sp.* regulatory sequence. **a**, Schematic of the CASK transcriptional reporter and eight tiling mutations between the Csx34 binding motif and the ribosome binding site. For each mutation, 39 bp of endogenous sequence was substituted with a control sequence to preserve spacing. **b,** Luciferase activity is greatly reduced in tiling mutants R3– R5. **c,** Schematic and sequence of a predicted promoter (P1), 153 bp downstream of the Csx34 binding motif and two additional putative promoter sequences (P2, P3). For each promoter, substitution mutations were made in the –35 and –10 elements. **d,** Luciferase activity of promoter mutants identifies P1 as a key element for expression of the native CASK operon. **e,** Csx33–Csx34 T269E upregulates luciferase activity even in a promoter P1 mutant reporter. **f,** Schematic of a CASK reporter replacing the P1 promoter with the T7 promoter. **g,** Luciferase activity in cells containing a T7 promoter and a plasmid expressing T7 phage polymerase confirming orthogonal expression. **h,** Luciferase activity in cells expressing Csx33–Csx34 with the T7 promoter reporter in the presence (red) or absence (blue) of T7 polymerase. In (e) and (h), expression was normalized to cells carrying an empty control plasmid. In (b), (d), (e), (g), and (h), error bars represent the standard deviation from the mean. *n* = 3–6 replicates.

**Extended Data Figure 14.**
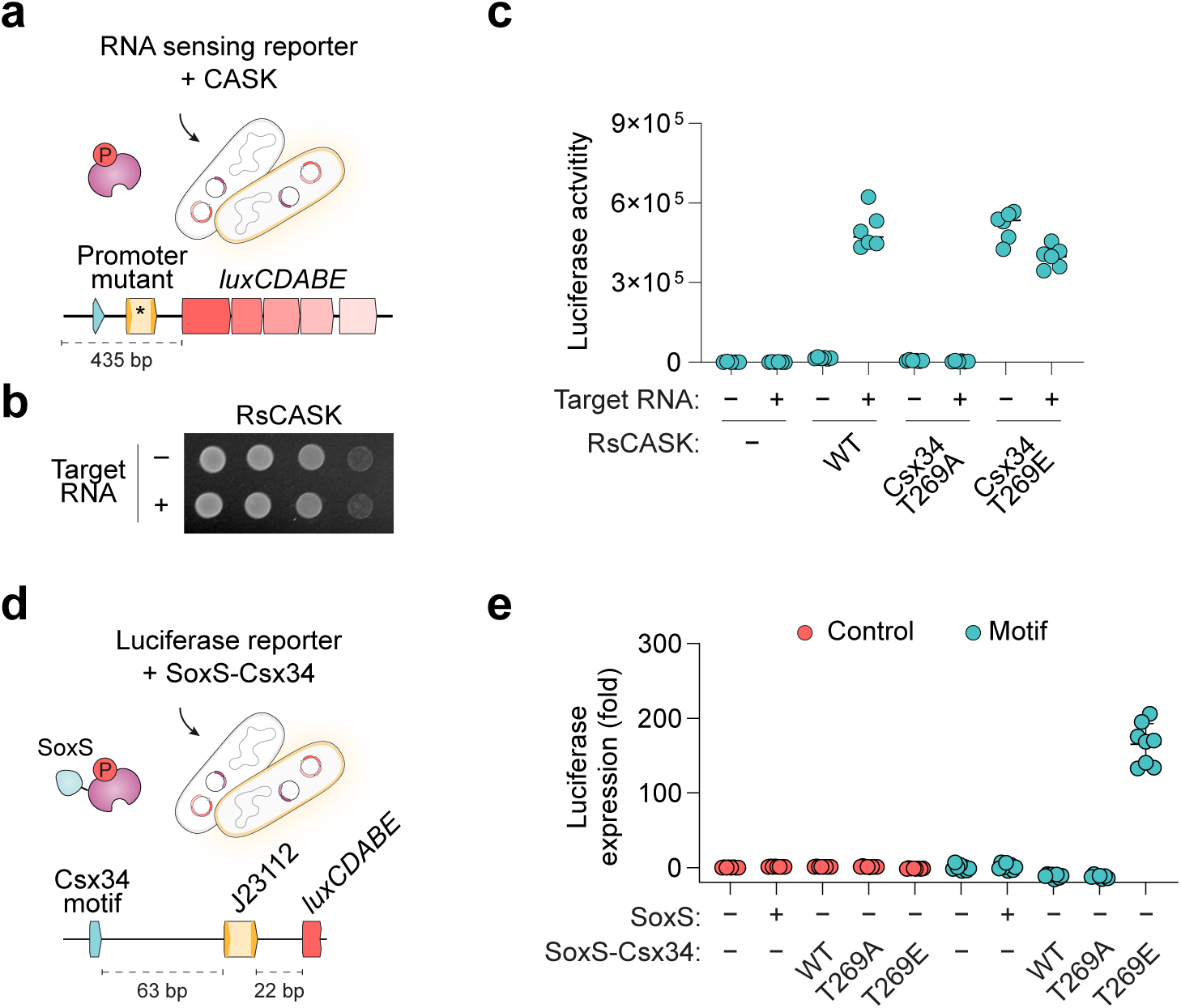
RNA sensing with RsCASK in *E. coli*. **a**, Schematic of a CASK transcriptional reporter containing a promoter P1 mutation. **b,** Viability of cells expressing CASK components in the presence or absence of target RNA under low induction conditions. Cells were serially diluted tenfold before spotting. **c,** RNA sensing of a target RNA in *E. coli* expressing RsCASK and a targeting (+) or non-targeting (–) crRNA. **d,** Schematic of a synthetic reporter containing the Csx34 binding motif upstream of the J23112 promoter in SoxS–Csx34-expressing cells. **e,** SoxS-mediated gene expression is dependent on Csx34 T269E and the binding motif sequence. Expression was normalized to cells carrying an empty control plasmid. In (c) and (e), error bars represent the standard deviation from the mean. *n* = 6–8 replicates.

**Extended Data Figure 15.**
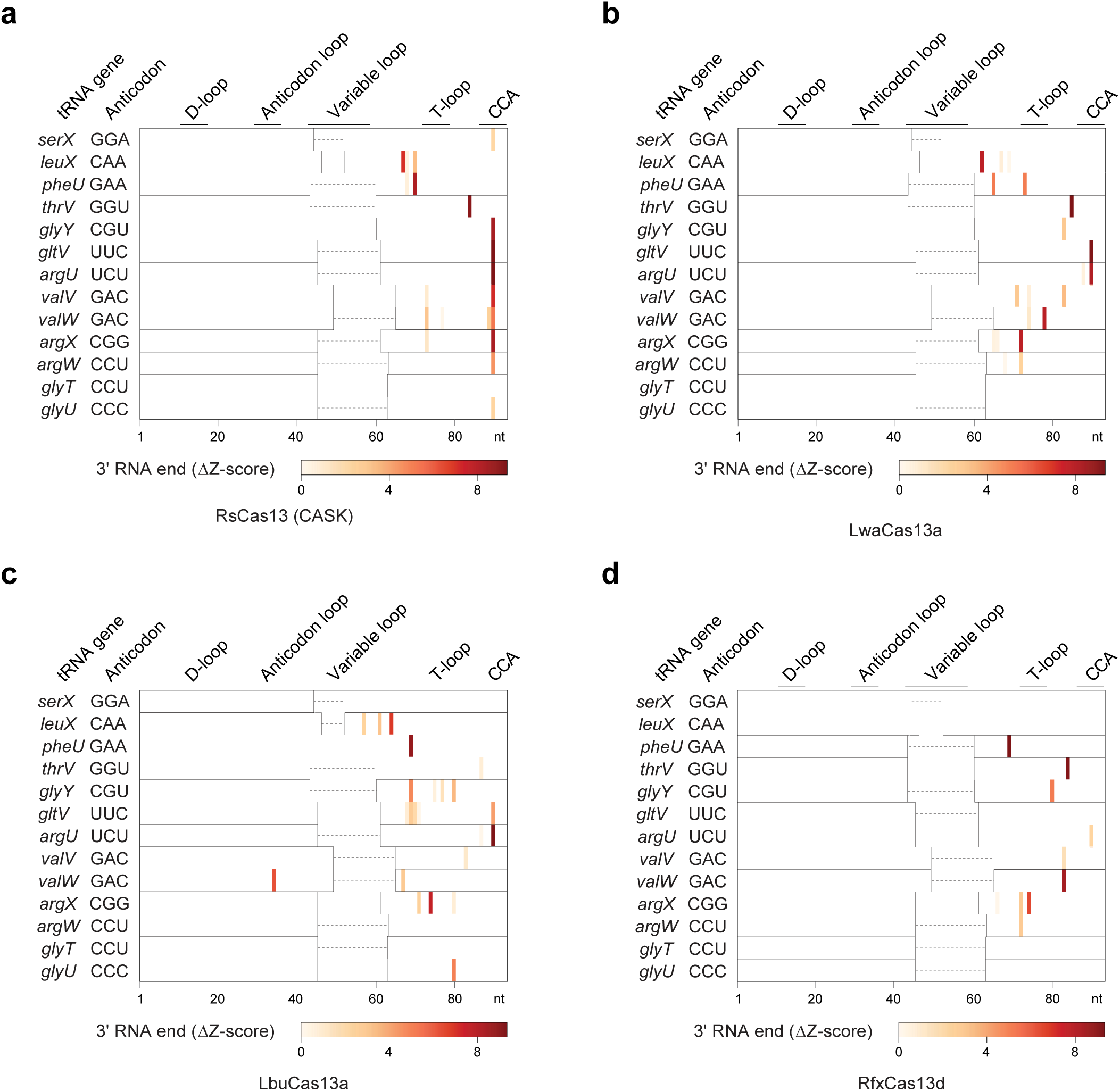
Collateral activity of diverse Cas13 orthologs on tRNA. **a–d**, In vitro Cas13 collateral reactions containing a pool of in vitro–transcribed *E. coli* tRNAs. Novel 3′ tRNA ends were detected by RNA sequencing following in vitro reactions containing target RNA, compared to a non-target RNA control. Reactions were performed with RsCas13 (a), LwaCas13a (b), LbuCas13a (c), and RfxCas13d (d).

